# A Multiverse Approach to Brain Age Prediction: Exploring the Impact of Modality, Bias Adjustment, and Atlas

**DOI:** 10.64898/2026.10.12.758514

**Authors:** J. Festor, L. Smolders, W. De Baene

## Abstract

Brain age, an individual’s age predicted from neuroimaging data, is a promising biomarker of brain health. Yet brain age estimates may depend on methodological choices within the prediction pipeline. We used a multiverse approach to systematically investigate effects of data modality, bias-adjustment method, and brain parcellation on brain-age prediction using data from 560 healthy adults (18.6-89 years) from the CamCAN dataset. Across 84 pipeline conditions, chronological age was predicted from connectivity features with elastic net regression, and performance was evaluated using mean absolute error (MAE). A multilevel model tested the effects of the pipeline factors and their interactions; moreover, cross-atlas consistency of the most predictive connections was assessed. MAEs varied substantially across conditions (5.26-18.30 years). Modality and bias adjustment had significant effects on MAE and interacted significantly. Concatenating structural and functional connectivity performed similarly to structural connectivity alone, and structure-function coupling produced the highest MAE in most conditions. Beheshti-based bias adjustment outperformed Cole-based adjustment, while combining adjustment procedures did not consistently improve performance. Atlas choice had no main effect on MAE, its interaction with modality was significant but negligible in size, and finer parcellations did not systematically improve performance. However, the network-level features driving predictions showed limited spatial overlap across atlases, and importance was distributed over many features. Brain-age prediction is thus sensitive to methodological choices, particularly modality and bias-adjustment strategy. Atlas choice appears less important for predictive accuracy but may affect which connections appear most predictive. These findings highlight the importance of systematically evaluating analytical choices in brain-age research.

---

As societies age and life expectancy increases, age-related diseases are becoming more prevalent, placing a growing burden on individuals and healthcare systems alike (Department of Economic and Social Affairs United Nations, 2023; World Health Organization, 2025). Improving early detection, prognosis, and treatment of these conditions is therefore of great importance in medical research. Central to these efforts is the discovery and development of reliable biomarkers, defined as objectively measurable indicators of normal or pathological biological processes (Manera et al., 2023; Strimbu & Tavel, 2010).

One promising biomarker is *brain age*, defined as an estimate of an individual’s biological age derived from neuroimaging data (Baecker et al., 2021; Franke et al., 2010; Franke & Gaser, 2019). Brain age is most informative when evaluated relative to chronological age. The difference between predicted age and chronological age is referred to as the brain age gap (BAG) (Franke et al., 2010; Franke & Gaser, 2019; Jirsaraie, Gorelik, et al., 2023; Kumari & Sundarrajan, 2024; Salih et al., 2023), which is an index of deviation from typical aging trajectories. A positive BAG (i.e., predicted age exceeding chronological age) is interpreted as an indicator of accelerated brain aging, whereas a negative BAG (i.e., chronological age exceeding predicted age) indicates delayed aging (Franke & Gaser, 2019). Depending on the context, variation in aging speed might represent a risk or protective factor for an individual’s health (Franke & Gaser, 2019).

To date, the brain age gap has been related to individual variation across a broad range of domains, spanning psychopathology (e.g., posttraumatic stress disorder, schizophrenia, depression, psychosis, obsessive-compulsive disorder, autism) (Cohen et al., 2024; Franke & Gaser, 2019; Jirsaraie, Gorelik, et al., 2023), socioeconomic indicators (e.g., household income, parent occupational prestige, years of education) (Cohen et al., 2024; Jawinski et al., 2022), neurodegenerative and medical conditions (e.g., Alzheimer’s disease, HIV, traumatic brain injury, type 2 diabetes) (Franke & Gaser, 2019; Jirsaraie, Gorelik, et al., 2023), genotypes (e.g., APOE) (Franke & Gaser, 2019), biophysiological markers (e.g., blood pressure, fasting blood glucose) (Jawinski et al., 2022; Jirsaraie, Gorelik, et al., 2023), lifestyle factors (e.g., physical activity, drug and alcohol use) and cognition (e.g., episodic memory, fluid intelligence) (Jawinski et al., 2022).

Because the brain age gap appears to reflect brain integrity and health (de Lange et al., 2022), brain age prediction has been proposed as a non-invasive and relatively cost-efficient tool (Kumari & Sundarrajan, 2024) to assist in early detection and prognosis of disorders, support differential diagnosis, and evaluate treatment outcomes (Baecker et al., 2021).

Despite growing interest, the impact of key methodological decisions in the brain age prediction pipeline remains poorly understood. Choices regarding input modality, bias-adjustment techniques, and brain parcellation scheme may substantially influence predictive performance and downstream interpretations of the BAG. In the following sections, we summarize the rationale for a systematic evaluation of these three dimensions.

With regard to data modality, structural features, whether derived from T1-weighted images (e.g., grey matter volume) or from diffusion-weighted images (e.g., tractography-based connectivity), have been used far more often in brain age prediction than functional indicators, and early models relied primarily on T1-weighted structural MRI features (Franke et al., 2010; Franke & Gaser, 2019). Functional indicators of neural activity (e.g., blood-oxygen-level-dependent signals) have also been shown to yield adequate brain age predictions, although their adequacy and predictive performance largely depend on the phenotype of interest (Jirsaraie, Kaufmann, et al., 2023): Structural features are often considered strong predictors because of their relative stability and strong associations with chronological age, while functional features may be more sensitive to transient or early pathological processes (Jirsaraie, Gorelik, et al., 2023). One important limitation in the current literature is that structural and functional modalities are typically examined in isolation (Corps & Rekik, 2019; Jirsaraie, Gorelik, et al., 2023; Kumari & Sundarrajan, 2024), despite the fact that the brain’s structural architecture and functional dynamics are intrinsically intertwined and dependent (Fotiadis et al., 2024). Integrating modalities may therefore capture complementary aspects of neurobiological aging and enhance predictive performance (Jirsaraie, Gorelik, et al., 2023). Here, we focus on connectivity-based features: structural connectivity (SC), derived from tractography on diffusion-weighted MRI, and functional connectivity (FC), derived from resting-state fMRI. One straightforward integration strategy involves concatenating SC and FC features into one single dataset. A more biologically informed alternative is using structure-function coupling (SFC), which quantifies how well a brain region’s structural connectivity matches its functional connectivity. SFC is typically operationalized as the correlation between a region’s SC and FC profiles (Fotiadis et al., 2024). Although promising, such coupling metrics have rarely been incorporated into brain age modelling frameworks.

A second methodological challenge concerns a bias in brain age estimations. Linear predictions commonly exhibit a systemic regression-to-the-mean effect in which older individuals tend to be predicted as younger while younger individuals are predicted to be older. This phenomenon reflects a statistical artefact of the chosen regression methodology rather than true individual variation (de Lange et al., 2022). To address this issue, several correction approaches have been proposed. Sample-level bias adjustment methods reduce the global bias by using the slope and intercept of a fitted regression line. Two widely adopted variations are the Cole and Beheshti adjustments (Zhang et al., 2023). Among these, Beheshti-based adjustments have been shown to better retain predictive performance, since the age-dependent offset is subtracted from each prediction rather than rescaling the predictions by dividing by the regression slope. Nevertheless, evidence suggests that such global sample-level corrections may not fully eliminate the bias: residual bias may persist within individual age strata (e.g., all participants aged 24) even when the overall sample bias appears corrected (Zhang et al., 2023). Consequently, an age-level bias adjustment method has been proposed to address systematic deviations within each age group in addition to the full sample (Zhang et al., 2023). Finally, Niu and colleagues (2019) proposed a nonlinear bias adjustment approach to account for the fact that age-dependent bias may follow nonlinear patterns not fully captured by linear corrections alone. Because these bias-correction procedures directly modify model predictions, they can inadvertently affect the strength or direction of associations between the brain age gap and cognitive or clinical outcomes. Yet, their relative impact on the predictive performance has not been systematically compared.

A final important methodological consideration concerns the choice of the brain atlas used to define regions and connections. Atlases vary in anatomical basis, functional assumptions and granularity, thereby shaping both feature reconstruction and neurobiological interpretation. Prior research has demonstrated that parcellation choice can substantially alter findings on connectivity and associations with age, poverty, or cognitive function (Bryce et al., 2021). Atlas selection can also change which brain regions appear to contribute most to age prediction, complicating neurobiological interpretation and reproducibility across studies. Evaluating the robustness of findings across multiple atlases is therefore essential for reliable inference.

Taken together, brain age research depends heavily on analytical choices that are to a large extent arbitrary, so any single methodological pipeline may not capture true neural aging processes. Multiverse studies address this issue by systematically evaluating how alternative methodological decisions affect outcomes. Prior multiverse work in brain age varied preprocessing and feature extraction choices across functional connectivity pipelines (Dafflon et al., 2022) or systematically compared multiple workflows combining different structural feature representations and machine learning algorithms (More et al., 2023). Notably, these studies focused on a single modality and did not systematically vary bias-adjustment strategies or brain parcellation schemes.

The present study extends prior work by manipulating three analytical dimensions (i.e., data modality, bias-adjustment technique, and atlas selection) within a single multiverse framework, thereby enabling a more comprehensive evaluation of how interdependent pipeline decisions jointly shape brain age predictions. By identifying which methodological choices strongly influence predictive performance, the study aims to provide concrete, evidence-based guidance to greater transparency and reproducibility in brain age research.

This study addresses four research questions. First, the study examines whether integrating structural and functional connectivity improves predictive performance relative to unimodal models (RQ1). It is expected that multimodal models combining SC and FC will outperform unimodal models (H1.1) because the two modalities capture complementary aspects of neurobiological aging (Franke & Gaser, 2019; Jirsaraie, Kaufmann, et al., 2023; Niu et al., 2019). In addition, models based on structure-function coupling are expected to outperform the concatenated multimodal model (H1.2) due to their more integrated representation of brain organization.

Second, the study investigates how different bias-adjustment methods applied to brain age estimates affect prediction performance (RQ2). It is expected that performance will monotonically increase as more bias adjustment methods are combined (i.e., sample-level adjustment, age-level adjustment, and nonlinear bias correction; H2.1). This premise is grounded in the belief that additional bias adjustment should reduce error variance. Furthermore, it is anticipated that Beheshti sample-level adjustments will yield more accurate brain age estimates than the Cole approach (H2.2), consistent with prior findings (Zhang et al., 2023).

Third, the study evaluates the sensitivity of brain age predictions to atlas selection (RQ3). It is expected that finer-grained parcellation schemes will yield lower prediction error because they capture more spatially specific aging patterns (H3). More broadly, substantial variability across atlases is expected, in line with prior evidence that parcellation choice can meaningfully alter connectivity findings (Bryce et al., 2021).

Finally, the study assesses whether the network-to-network connections identified as most predictive of brain age remain consistent across parcellation schemes (RQ4). Based on previous findings demonstrating only moderate overlap between networks derived from different atlases in healthy aging (Deschwanden et al., 2024), the specific connections identified as most predictive of brain age are expected to vary across atlases (H4).

To address these questions, the present study used the Cambridge Centre for Ageing and Neuroscience (CamCAN) dataset (Shafto et al., 2014), selected for its broad adult lifespan coverage (18-88) and the availability of both diffusion-weighted and resting-state fMRI data. A multiverse consisting of 84 pipeline conditions was constructed by systematically varying modality, bias-adjustment method, and parcellation scheme. Machine learning was used to generate brain age predictions across all conditions, while the consistency of regional contributors across parcellation frameworks was assessed by mapping regions onto broader functional networks.

## Methods

### Dataset

Data used in the preparation of this work were obtained from the CamCAN repository (Shafto et al., 2014; available at https://www.mrc-cbu.cam.ac.uk/datasets/camcan/). The dataset (Phase 2, Arm1) initially contained 652 participants with resting-state fMRI data and 641 with diffusion-weighted imaging data. Participants with complete data across all three image measures (a 3D T1-weighted structural image, diffusion-weighted images (DWI), and resting-state T2*-weighted fMRI) were retained, yielding a sample of 641 healthy adults.

Ethical approval as part of the CamCAN obtainment was granted by the Cambridgeshire 2 Research Ethics Committee (reference: 10/H0308/50), in compliance with the Helsinki Declaration (Shafto et al., 2014).

### Workflow

The present study followed a multi-stage pipeline, progressing from preprocessing through feature extraction, age prediction, and bias adjustment prior to statistical testing. The following sections describe each step in detail. Figure 1 provides a schematic overview of the workflow.

**Figure 1.**
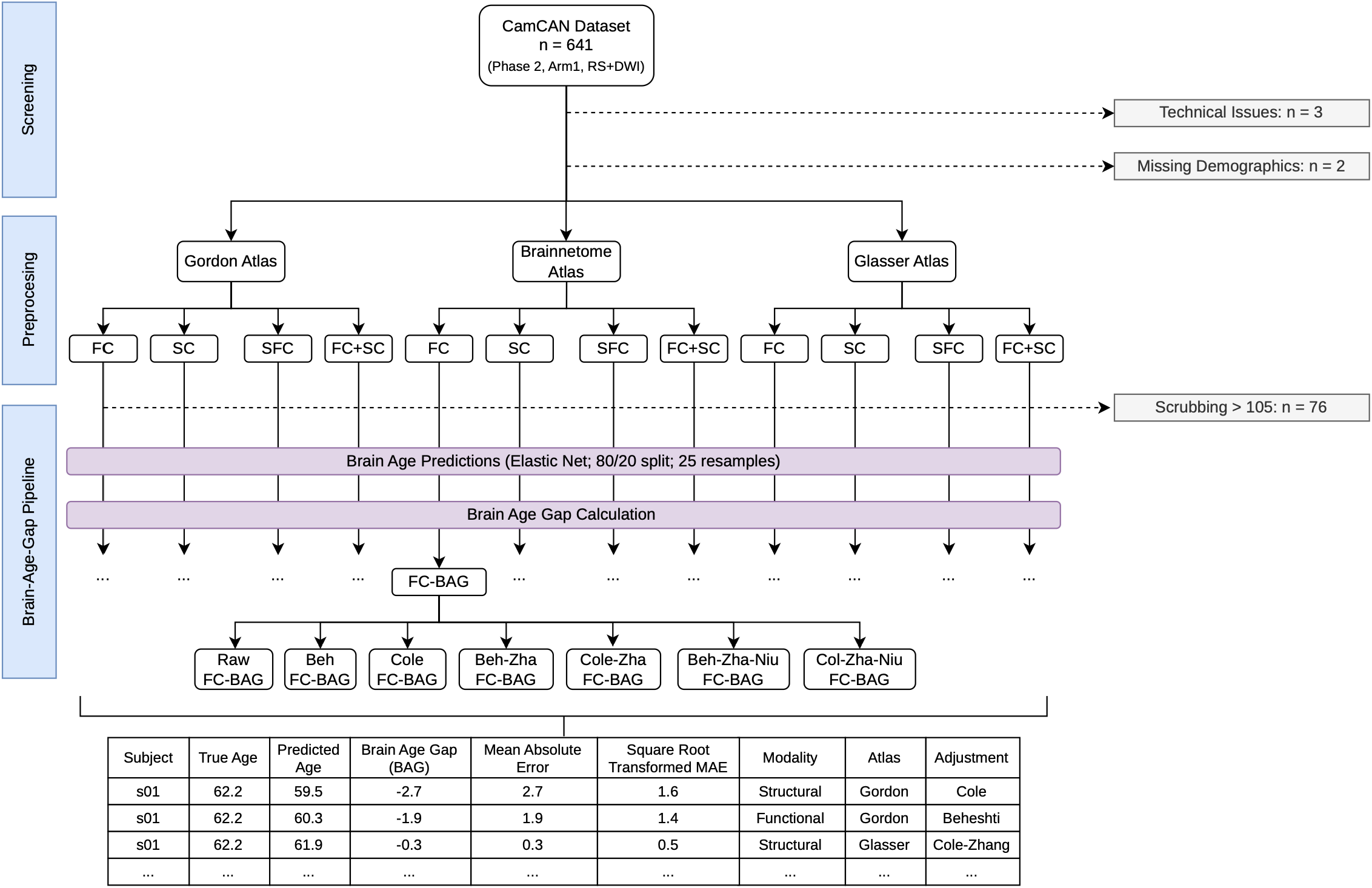
Brain Age Prediction Workflow: From Screening to Estimation. *Note*: Dotted arrows = exclusions; for each branch the same participants (n = 560) are used; FC = functional connectivity, SC = structural connectivity, SFC = structure-function coupling, FC+SC = concatenated functional and structural connectivity; Beh = Beheshti adjustment, Zha = Zhang adjustment; BAG = brain age gap; RS+DWI = sample of participants featuring resting state and diffusion weighted images; output table based on fictional data for illustrative purpose only.

### Measures

#### Image Acquisition

The image acquisition steps followed the protocol described by Shafto et al. (2014). Imaging was performed on a 3T Siemens TIM MRI scanner equipped with a 32-channel radiofrequency head coil. The T1-weighted structural images were acquired using a Magnetization Prepared Rapid Gradient Echo (MPRAGE) sequence (TR/TE/TI = 2250/2.99/900ms; FOV = 256 x 240 x 192mm; voxel size = 1 mm isotropic). The diffusion-weighted images were acquired using a twice-refocused spin-echo echo-planar imaging sequence (TR/TE = 9100/104ms; FOV = 192 x 192 mm; voxel size = 2mm isotropic) comprising 30 directions at b = 1000 s/mm^2^, 30 directions at b = 2000 s/mm^2^ and 3 images with b = 0 s/mm^2^. Resting state data were acquired using a Gradient-Echo Echo-Planar Imaging (EPI) sequence including 261 volumes with each 32 axial slices (TR/TE = 1970/30ms; FOV = 192 x 192 mm; voxel size = 3 x 3 x 4.44 mm; slice thickness = 3.7 mm).

#### Preprocessing

##### Resting State

Preprocessing of functional and structural MRI data was performed using the CONN toolbox version 22 (Nieto-Castanon & Whitfield-Gabrieli, 2022) and SPM12 (Friston et al., 2006), implemented in MATLAB version R2025a (The MathWorks, Inc., 2025). The preprocessing pipeline proceeded in the following stages. First, functional images were realigned with susceptibility distortion correction, followed by slice-timing correction and outlier detection using ART (framewise displacement threshold = 0.5 mm; global BOLD signal threshold = 3 SD). Next, structural and functional images were both segmented and normalized into standard MNI space and resampled to 2 mm isotropic voxels. Functional images were then spatially smoothed using a Gaussian kernel of 8 mm full width at half maximum (FWHM).

Functional timeseries were denoised using the standard CONN denoising pipeline. Sources of spurious variance were removed by linear regression of white matter and CSF signals estimated using the CompCor approach (5 components each), 6 motion parameters and their first-order derivatives (12 regressors in total), ART-derived outlier volumes, session effects, and linear trends. Finally, temporal bandpass filtering was applied between 0.01–0.1 Hz to retain frequencies associated with resting-state neural fluctuations.

##### Diffusion

Preprocessing of diffusion-weighted MRI images was performed using MRtrix3 (Tournier et al., 2019). Diffusion volumes were first denoised followed by correction for Gibbs ringing artifacts (Bautista et al., 2021; Kellner et al., 2016). Susceptibility-induced distortions, subject motion and eddy-current distortion were corrected (Andersson et al., 2003; Andersson & Sotiropoulos, 2016), and bias field correction was applied (Tustison et al., 2010). All DWI images were resampled to 1.5 mm isotropic voxel size. All preprocessing outputs were visually inspected for quality assurance.

T1-weighted anatomical images were skull-stripped using HD-BET (Isensee et al., 2019). To improve cross-modal registration accuracy, an inverted-contrast version of the skull-stripped T1 image was generated and nonlinearly registered to the mean b0 image using ANTs (https://github.com/ANTsX/ANTs; Avants et al., 2011). The resulting transforms were used to project anatomical segmentations and atlas parcellations into native diffusion space.

Orientation distribution functions (ODFs) were separately generated for white matter (WM), gray matter (GM) and cerebrospinal fluid (CSF) using multi-shell multi-tissue constrained spherical deconvolution (MSMT-CSD, Jeurissen et al., 2014), followed by global intensity normalization (Raffelt et al., 2017) to ensure cross-subject comparability.

Anatomically Constrained Tractography (ACT, Smith et al., 2012) was applied to improve the biological plausibility of reconstructed streamlines. The T1-weighted image registered to diffusion space was segmented into five tissue types (5TT), from which a GM/WM boundary mask was derived to guide streamline seeding. Whole-brain probabilistic tractography was performed using the iFOD2 algorithm as implemented in MRtrix3. Tracking was performed with ACT backtracking and GM-WM interface cropping enabled, an FOD cutoff of 0.1, minimum streamline length of 5 mm, maximum streamline length of 250mm, and generation of 5 million streamlines.

The tractogram was subsequently filtered using spherical-deconvolution informed filtering of tractograms (SIFT2; Smith et al., 2015), which assigns weights to individual streamlines in the whole-brain tractogram in proportion to their contribution to the intra-axonal diameter. Structural connectivity matrices were constructed by summing SIFT2-weighted streamlines between each pair of atlas-defined regions of interest using tck2connectome with a 4 mm search radius, scaled by the subject-specific SIFT2 fiber density proportionality coefficient (µ). Connectivity matrices were computed for all atlases described below after transforming them from MNI space into native diffusion space using ANTs.

#### Feature Extraction

Three widely used and neurobiologically distinct brain atlases were applied independently: the Human Brainnetome atlas, the Gordon atlas and the Human Connectome Project-Multimodal Parcellation atlas (HCP MMP1.0). The Human Brainnetome atlas (Fan et al., 2016) is a structural and functional connectivity-based parcellation comprising 246 cortical and subcortical regions. The Gordon atlas (Gordon et al., 2016) provides a parcellation of 333 cortical regions derived from resting-state functional connectivity. Finally, the HCP MMP1.0 atlas (Glasser et al., 2016) is a multimodal cortical parcellation combining structural, functional, and myelin-based features to define 360 regions.

For each atlas, parcellations were consistently applied across all imaging modalities, yielding the following four feature sets per atlas:

- Functional Connectivity (FC): pairwise FC between regions was computed as the Pearson correlation between regional BOLD timeseries, followed by Fisher z-transformation.
- Structural Connectivity (SC): SC was derived from diffusion MRI tractography and quantified using SIFT2-weighted streamline counts.
- Structure-Function Coupling (SFC): SFC was computed at the regional level as the Spearman correlation between a region’s full profile of structural and functional connections.
- Concatenated FC+SC: FC and SC matrices were combined into a single feature set, such that each participant is represented by both modalities.

#### Exclusions

Exclusion criteria were applied to the sample of 641 participants. Three participants were removed from all analyses due to technical issues in diffusion image preprocessing. Additionally, two participants were excluded due to missing demographic information. Seventy-six participants were excluded because they had more than 105 scrubbed timepoints (i.e., >40% of all timepoints) since excessive motion-related data loss compromises signal quality. These criteria resulted in a final sample of 560 participants.

### Statistical Analysis

All analyses, including brain age prediction, brain age gap transformation, bias adjustment, multilevel modelling, regional embedding, and additional exploratory analyses, were conducted in R (version 4.5.1; R Core Team, 2025) using Positron (version 2026.09.0; Posit Team, 2026).

#### Prediction Pipelines

Brain age prediction was operationalized as a supervised regression problem, in which a machine learning model was trained to predict chronological age from neuroimaging-derived features. The brain age gap was subsequently quantified as the mean absolute error (MAE), defined as the average of the absolute deviation between predicted and chronological age in years.

A preliminary analysis was conducted to identify the most suitable machine learning algorithm for the high-dimensional neuroimaging data. Specifically, functional connectivity data parcellated with the Gordon atlas (n = 560 participants) was used to compare candidate algorithms. Considering predictive performance, computational feasibility and interpretability, regularized regression (i.e., elastic net) was selected for all subsequent analyses (see supplementary material A, SM.A).

To ensure comparability across all analytical conditions and avoid variability induced by different train-test splits, a single central 80/20 train–test split was created and applied consistently across all pipelines. The split was stratified on chronological age to preserve the age distribution in both training and test sets. Since brain age predictions were carried out for each modality (four levels) × atlas (three levels) combination, twelve elastic net models were fit to the data predicting chronological age. Within each training set (determined by central test-train split), hyperparameters were tuned using five-fold cross-validation repeated five times (25 resamples). Regularization parameters (λ, α) were selected to minimize mean absolute error (MAE). Importantly, MAE served exclusively as the model selection criterion, whereas regression coefficients were estimated by minimizing the penalized least-squares objective (detail in SM.B).

All predictors were standardized independently (i.e., centered and scaled) across all subjects prior to model fitting. Variable importance was quantified as the absolute value of the standardized regression coefficients. Moreover, final age-predictions and performance metrics (i.e., R², MAE, RMSE) were computed on the held-out test set and retained for further analyses.

#### Bias-Adjustment Methods

Bias correction procedures were applied sequentially to the 12 elastic net test set predictions to adjust for systematic bias in linear age predictions. First, standalone Cole and Beheshti adjustments were implemented, each addressing sample-level bias (Zhang et al., 2023). Subsequently, the Zhang adjustment was applied on top of the Cole and Beheshti corrections to additionally account for age-specific linear bias (Zhang et al., 2023). Finally, the non-linear correction proposed by Niu et al. (2019) was applied to the Zhang–Cole and Zhang–Beheshti pipelines to additionally correct for non-linear age-related bias. Thus, in addition to the no-adjustment control condition, six bias-adjustment strategies were implemented. Crossing adjustment method (seven levels) with modality (four levels) and atlas (three levels) resulted in a total of 84 experimental conditions.

To prevent data leakage, all bias-adjustment parameters were estimated exclusively on out-of-fold (OOF) predictions from the training set and subsequently applied to the held-out test set. OOF predictions were obtained from models trained on subsets of the training data that explicitly excluded the participant being predicted, thereby capturing out-of-sample generalization rather than in-sample fits (details in SM.C).

#### Multilevel Modelling

To examine the impact of modality (4 levels), bias-adjustment method (7 levels), atlas (3 levels) and their interactions on prediction error, a multilevel linear model was employed. Modality, bias-adjustment method, and atlas were specified as fixed effects, including their interaction terms. Participants were modelled with a random intercept to account for repeated measurements across conditions within individuals (see SM.D). The BOBYQA optimizer (Powell, 2009) was specified for the multilevel model. BOBYQA is a derivative-free optimizer that is well-suited to the irregular likelihood surfaces common in mixed models; the primary motivation for lme4’s adoption of derivative-free methods (Bates et al., 2015).

The dependent variable was the individual-level absolute prediction error. Because absolute error is bounded at zero and exhibited strong positive skewness, normality assumptions were not satisfied. A square-root transformation was applied as this transformation is generally recommended when deviations from normality are moderate (Tabachnick & Fidell, 2013). All inferential statistics were conducted on the square-root-transformed scale. For readability, the transformation is not explicitly indicated in the presentation of the statistical results.

Main and interaction effects were summarized in ANOVA tables. Planned contrasts were added to investigate research question one through three (RQ1-3). Within each analysis, p-values were adjusted using the false discovery rate (FDR) procedure (Benjamini & Hochberg, 1995). A conservative significance threshold of α = .01 was adopted to reduce the likelihood of Type I error, acknowledging the corresponding increase in Type II error risk. Such conservative alpha thresholds are particularly important in multiverse studies where the large number of comparisons across analytical pipelines increases the risk of false positives. More specifically, the strict alpha threshold was chosen because within-test FDR correction alone does not account for the inflated Type I error introduced by conducting multiple distinct tests (RQ 1-3).

Because the present study relied on the CamCAN dataset (Shafto et al., 2014) which has a fixed sample size, conventional a priori power analysis was infeasible. However, statistical power remains informative in fixed-n designs, as it characterizes the probability of detecting effects across a range of plausible magnitudes. Power was therefore estimated across a grid of effect sizes and intraclass correlation coefficients (ICC), where ICC variation reflected best-to worst-case clustering scenarios. Given 111 individuals in the held-out test set per condition, the resulting power estimates are reported per main effect in the supplementary materials E (SM.E).

#### Brain Region Embedding

To examine the overlap in the connections contributing most strongly to brain age predictions across atlases (RQ4), both unimodal and concatenated multimodal models were projected onto a common reference framework using the less granular Yeo-7 network parcellation (Yeo et al., 2011). Specifically, regions from each atlas were assigned to one of the seven cortical networks using a winner-takes-all mapping approach, while subcortical regions were grouped into an additional eighth category.

Overlap across atlases was quantified using the SuperExactTest framework (Wang et al., 2015), which extends conventional pairwise overlap testing to multi-set intersections and therefore allows simultaneous evaluation of both pairwise and higher-order overlaps rather than restricting comparisons to two atlases at a time. Because the test assumes independence between elements, and this assumption was likely violated here (top-ranked connections probably have some dependencies), we particularly looked at the metric of fold enrichment (FE) over the test itself. Fold enrichment (FE) was computed as the ratio of observed to expected overlap, where FE > 1 indicates greater overlap than expected by chance. We evaluated the overlap among the top-10 connections of the nested Yeo-7 connection space. The analysis was not extended to top-25 or top-50 connections since these thresholds are too large relative to the available connection space. In the unimodal models which have only 36 possible network–network connections (including within-network connections), a top-25 would cover most of the space and a top-50 would exceed the number of available connections. In the concatenated models (72 possible connections), a top-50 would likewise cover most of the space. Such large thresholds would inflate the estimated overlap and reduce interpretability.

#### Exploratory Analyses

Beyond the primary analyses addressing RQ 1-4, exploratory post-hoc analyses were conducted to further interrogate patterns emerging from the results. These analyses were not specified a priori but were motivated by questions raised during interpretation.

##### Comparison Coupling and Concatenated Dataset

This exploratory analysis was aimed at clarifying whether performance differences between structure-function coupling (SFC, 246-360 features) and concatenated multimodal models (60,270-129,240 features) reflected (a) differences in feature dimensionality or (b) differences in feature information content (cf., RQ1). To disentangle these two factors, elastic net models were trained on randomly sampled subsets of the concatenated dataset with systematically increasing feature sizes. For each atlas, sampling started at a feature count equal to that of the corresponding SFC representation and increased by successive doubling up to that atlas’s full concatenated feature space. For each feature size, 50 repetitions with independent random feature samples were evaluated on the held-out test set. Hyperparameters for the repeated fitting of the concatenated dataset were selected using 5-fold cross-validation (5CV) ones for each feature-set size, rather than ones for each repetition to reduce computational burden. The cross-validated SFC performance on the corresponding full feature set served as a reference baseline. This comparison therefore provides a slight advantage for the SFC, which benefited from model-specific hyperparameter optimization whereas the concatenated models used the same cross-validated hyperparameters across the 50 random feature samples at a given feature size. The concatenated condition has thus less robust hyperparameter tuning than the coupling data making it somewhat more difficult for the concatenated condition to outperform.

Prior to this exploration, variance homogeneity between edge-level (concatenated) and node-level (SFC) features was assessed separately per atlas to rule out the possibility that random feature sampling would be systematically biased by differences in feature variability. The median feature variance did not differ significantly. Thus, this confound was ruled out prior to exploration.

##### Cumulative Feature Importance in Atlas Comparison

To further characterize the cross-atlas overlap in predictive features (cf., RQ4), additional exploratory analyses examined the distribution of feature importance across atlases. Specifically, matrix sparsity was extracted for each atlas, and cumulative importance curves were constructed. The latter were quantified as the non-zero ranked features given their absolute beta value. These indicators are used to understand whether predictions were driven primarily by a small subset of highly predictive connections or distributed more broadly across the feature space.

## Results

### Descriptives

The final sample comprised 560 healthy adults with a mean age of 53.64 years (SD_age_ = 18.53, range: 18.6–89.0 years), of whom approximately half were female (48.9%). Individuals’ education varied (range: 9.0-18.0 years), however the distribution was skewed towards higher education (M_Education_ = 14.93, SD_Education_ = 2.73).

Brain age predictions were generated for this sample across 84 conditions (adjustment method (7) x modality (4) x atlas (3)). The performance of the brain age predictions ranged between a mean absolute error (MAE) of 5.26 and 18.30 years compared to chronological age (see Figure 2 & 3). The mean error was situated at 8.39 years (SD_MAE_ = 3.12). Explained variance ranged between 44.97 and 90.90%, with a mean of 77.1% (SD_R2_ = 13.35%; see SM.F). The *Nui-Zhang-Beheshti-adjusted Gordon pipeline with concatenated structural and functional features* achieved the lowest MAE and explained 90.08% of the variance in age (MAE = 5.26, RMSE = 6.58).

**Figure 2.**
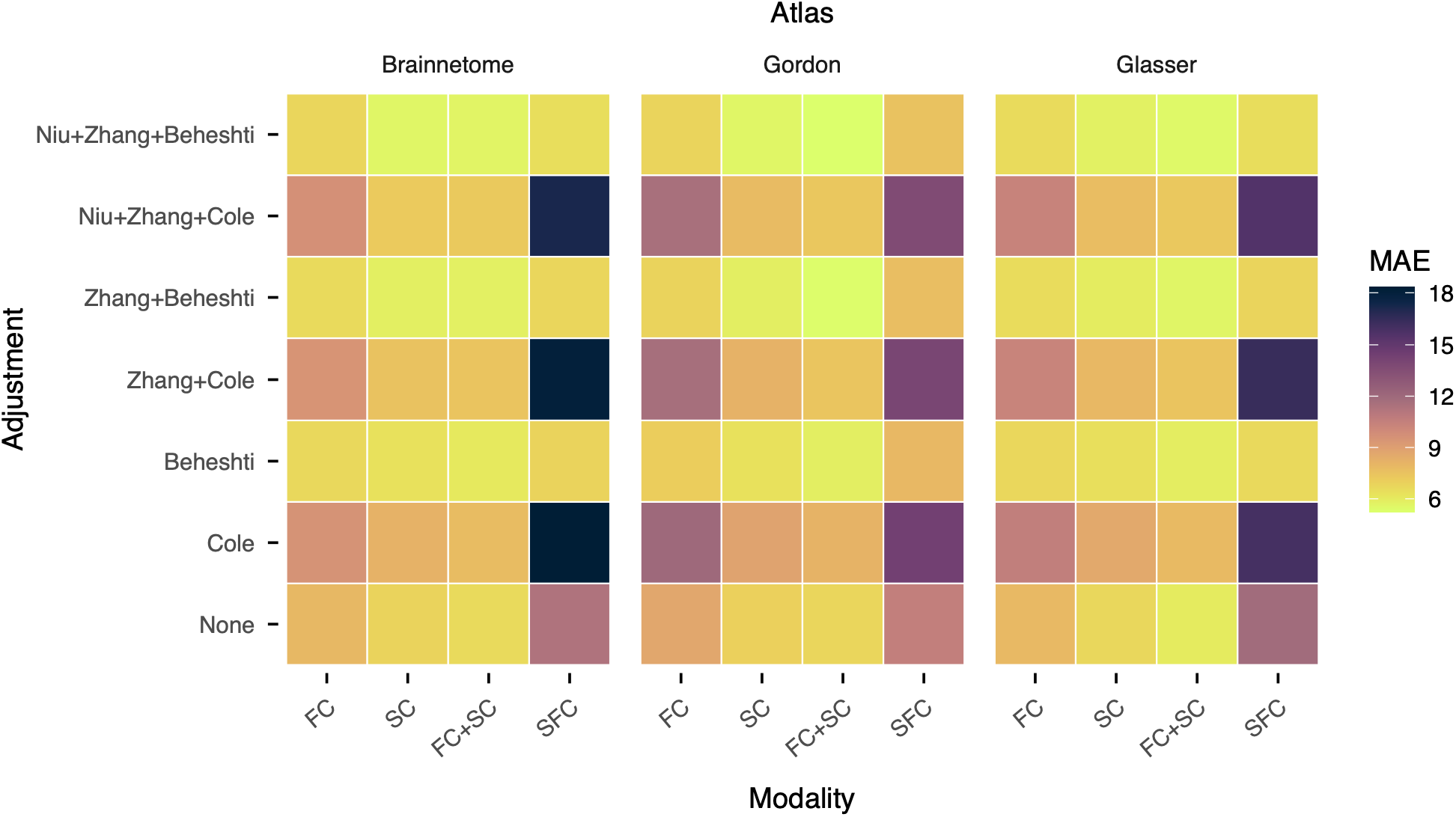
Heat-Map of Mean Absolute Error (MAE) Across 84 Experimental Conditions. *Note:* FC = functional connectivity, SC = structural connectivity, FC+SC = concatenated structural and functional connectivity, SFC = structure-function coupling.

**Figure 3.**
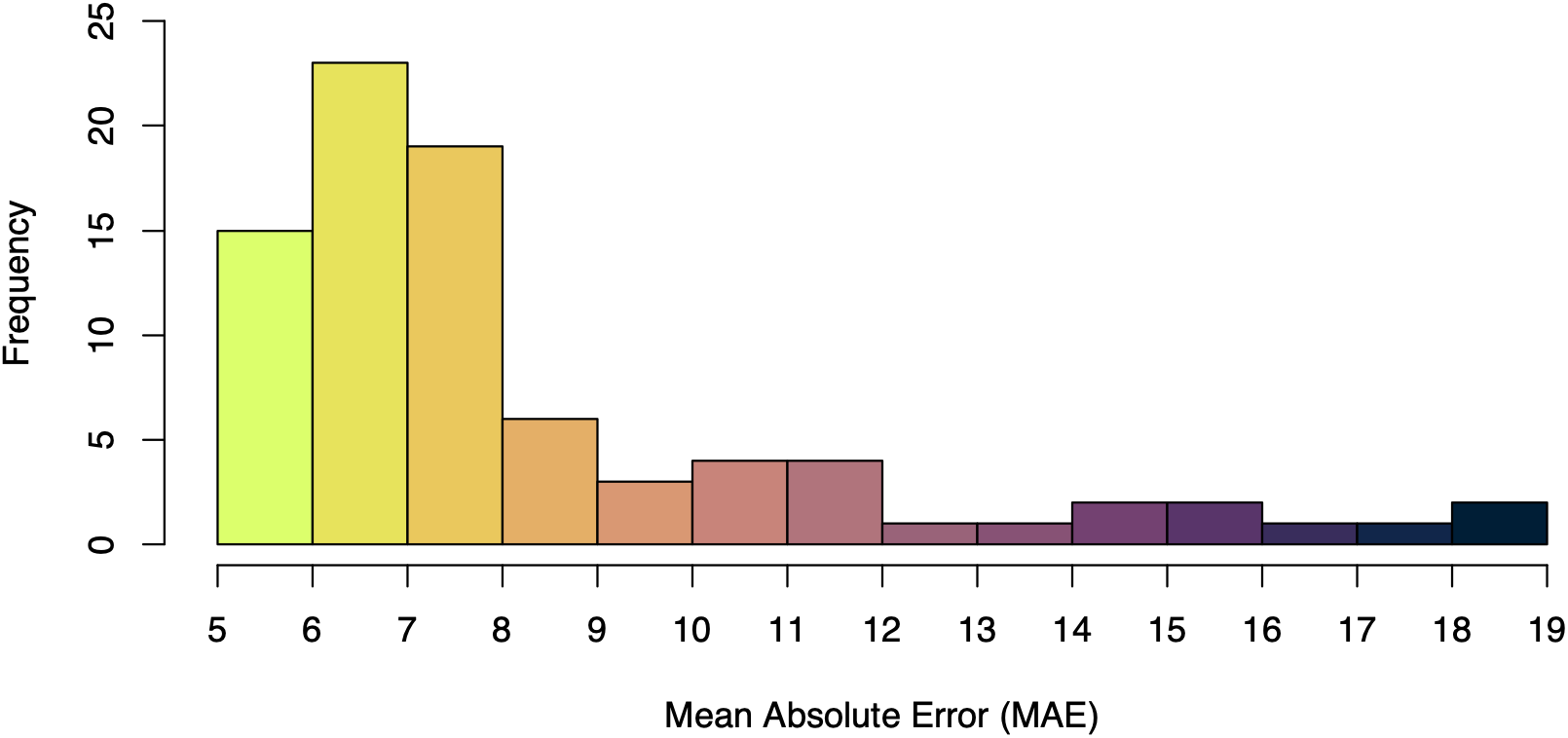
Frequency Distribution of Mean Absolute Error (MAE) Across Conditions. *Note:* Colouring represents the magnitude of mean absolute error (MAE) analogous to Figure 2 (darker colours = higher mean prediction error of age).

### Multilevel Model

The multilevel model with random intercept per participant revealed a main effect of Modality, *p(BH)* < .001, *η_p_^2^* = .070, and of Adjustment, *p(BH)* < .001, *η_p_^2^* = .080, but not of Atlas (see Table 2). Two two-way interactions were also significant: Modality x Atlas, *p(BH)* < .001, *η_p_^2^* = .004, and Modality x Adjustment, *p(BH)* < .001, *η_p_^2^* = .030. The ICC of the multilevel model was .208, indicating that 20.80% of variance in predictive performance (MAE) was attributable to differences between subjects. Conversely, 79.2% reflected within-subject changes, reflecting a combination of effects from the experimental-conditions and residual variability. Given the significant interactions, simple effects analyses were performed to answer the research questions in context of the interaction partners. Within each simple effects analysis, planned (a priori) contrasts were specified to test each hypothesis.

**Table 2.** Multilevel Model Results with Random Intercept per Participant.

| Effects | DFn | DFd | F | p(BH) | $\eta_p^2$ |
| --- | --- | --- | --- | --- | --- |
| Modality | 3 | 9130 | 224.15 | < .001 | .070 |
| Atlas | 2 | 9130 | 0.05 | .992 | < .001 |
| Adjustment | 6 | 9130 | 130.84 | < .001 | .080 |
| Modality * Atlas | 6 | 9130 | 6.32 | < .001 | .004 |
| Modality * Adjustment | 18 | 9130 | 14.43 | < .001 | .030 |
| Atlas * Adjustment | 12 | 9130 | 0.29 | .992 | < .001 |
| Modality * Atlas * Adjustment | 36 | 9130 | 1.36 | .106 | .005 |

*Note:* DFn = numerator degrees of freedom, DFd = denominator degrees of freedom, p(BH) = Benjamini-Hochberg adjusted p-values, $\eta_p^2$ = partial eta squared; dependent variable = square root transformed mean absolute error.

#### Effects of Modality (RQ1)

*Modality x Adjustment.* To test H1.1 and H1.2, we examined the simple effects of Modality within each adjustment condition. Reported effects reflect conditional main effects, averaged across atlases.

Contrary to H1.1, multimodal models (concatenated SC+FC and SFC) showed significantly higher MAE than unimodal models (SC and FC) for the majority of adjustment conditions (all ps < .001). This did not hold for Beheshti-based adjustments, namely, Beheshti only (p = .705), Zhang-Beheshti (p = .860) and Niu-Zhang-Beheshti (p = .661). Contrary to H1.2, SFC showed higher MAE than concatenated multimodal models across all adjustment conditions (all ps < .01), except for Beheshti only (p = .014).

To better understand the unimodal vs multimodal results (H1.1) (as visually depicted in Figure 4), we conducted follow-up contrasts comparing the concatenated multimodal model separately with FC and with SC. The concatenated model led to significantly lower MAE than FC across Cole, Zhang-Cole, Niu-Zhang-Cole, Niu-Zhang-Beheshti and no-adjustment conditions (ps < .01); however, no significance was reached for the Beheshti and Zhang-Beheshti condition (ps > .042). SC did not differ significantly from the concatenated model in any adjustment condition (all ps > .470).

**Figure 4.**
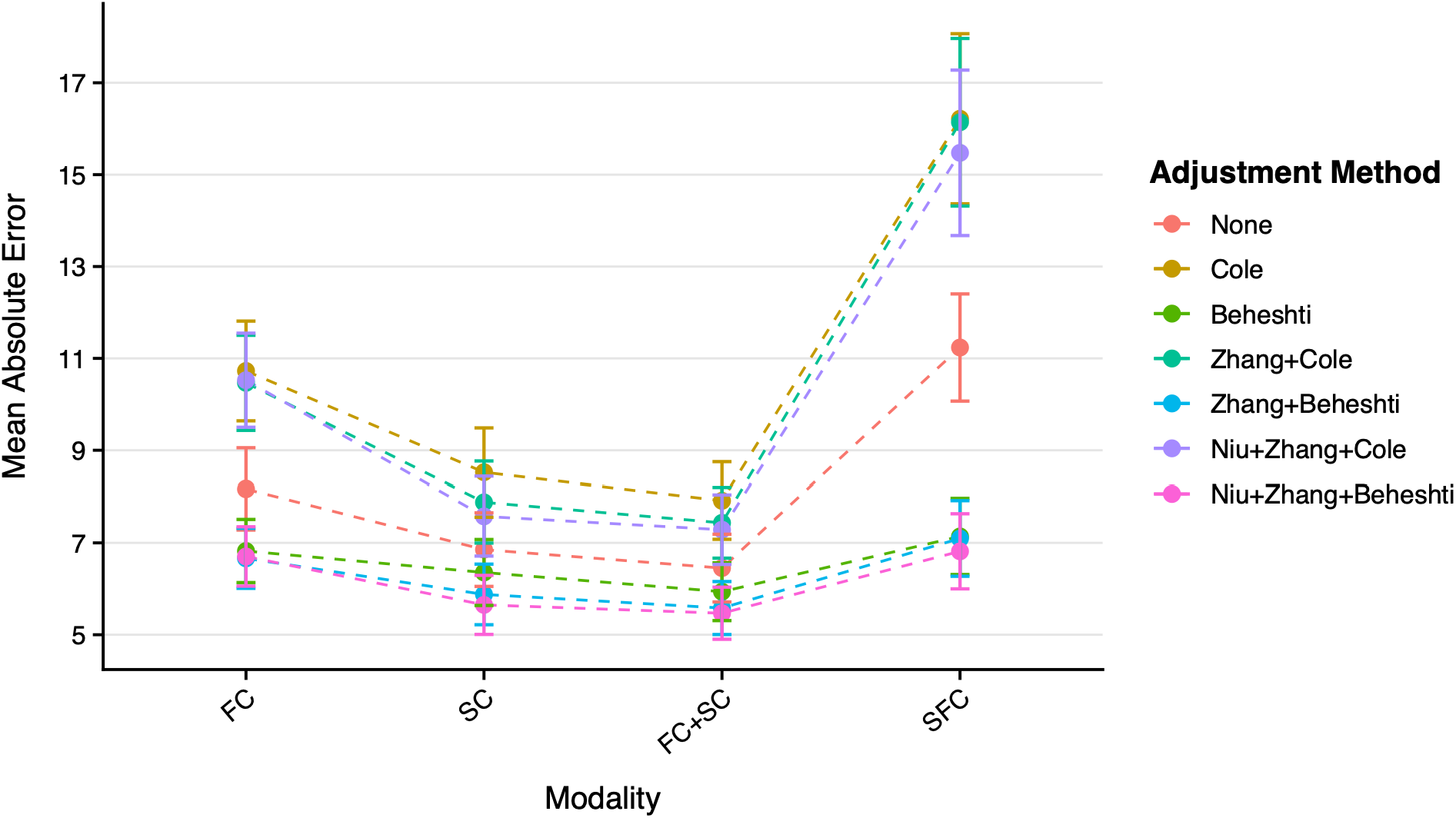
Interaction Plot: Mean Absolute Error by Modality x Adjustment. *Note:* 99% confidence interval used, calculated from raw absolute error for readability (not model-estimated, nor transformed data); FC = functional connectivity, SC = structural connectivity, FC+SC = concatenated structural and functional connectivity, SFC = structure-function coupling.

#### Modality x Atlas

A parallel analysis was conducted for the modality × atlas interaction, with effects averaged across adjustment method. Contrary to H1.1, age predictions from multimodal models led to significantly higher MAE than unimodal models for the Brainnetome and HCP MMP1.0 parcellations (all ps < .001) but not for the Gordon parcellation (p = .796). Additionally, SFC consistently showed higher MAE than the concatenated multimodal models across all atlas types (ps < .001, H1.2). Again, follow-up contrasts were conducted to clarify the unimodal vs multimodal results (H1.1)(visually depicted in Figure 5). The concatenated model resulted in significantly lower MAE than FC across all parcellations (ps < .001). However, the concatenated model and SC were not significantly different across parcellations (all ps > .219).

**Figure 5.**
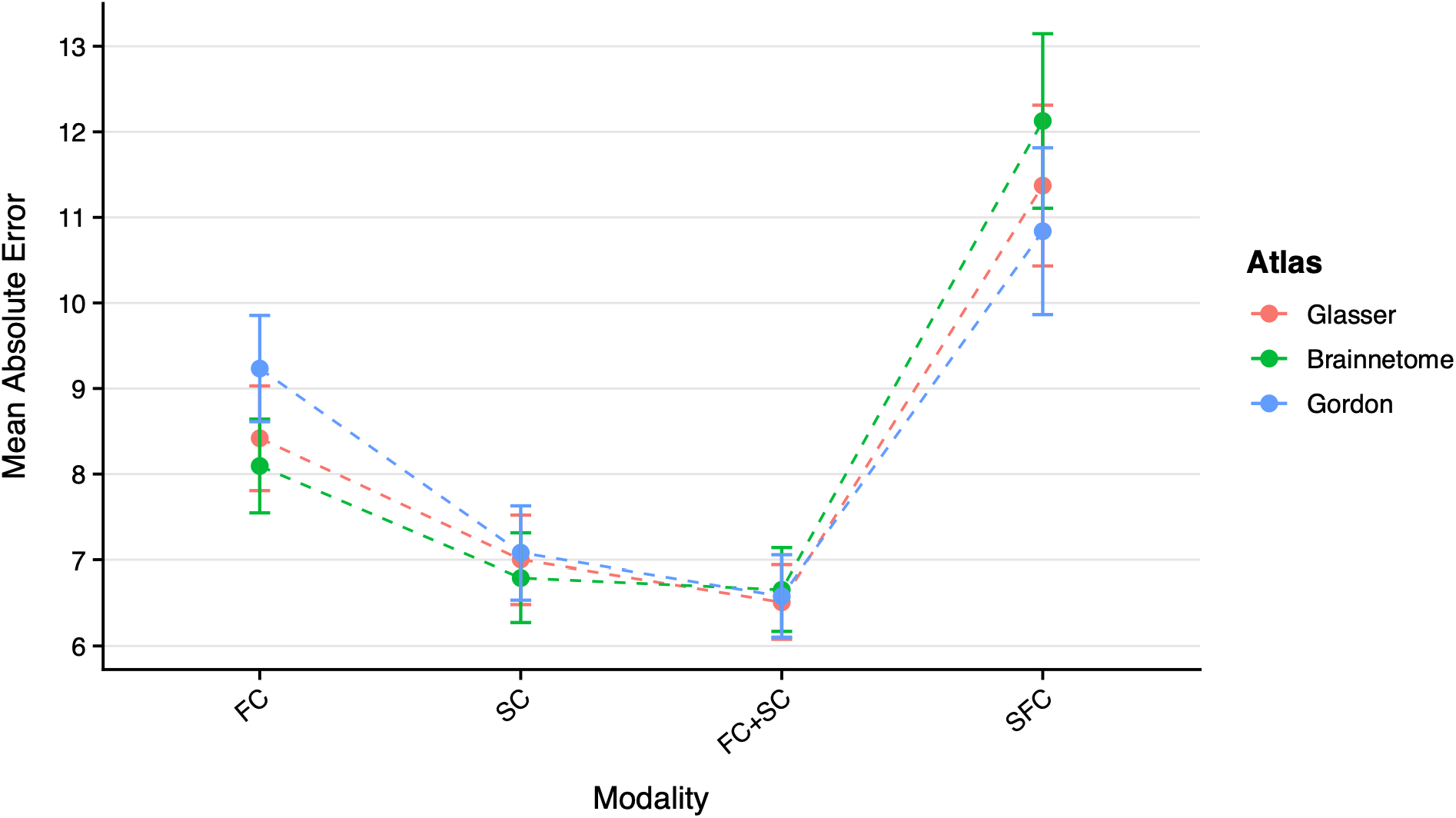
Interaction Plot: Mean Absolute Error by Modality x Atlas. *Note:* 99% confidence interval used, calculated from raw absolute error for readability (not model-estimated, nor transformed data); FC = functional connectivity, SC = structural connectivity, FC+SC = concatenated structural and functional connectivity, SFC = structure-function coupling.

### Exploratory Analysis: Coupling vs. Concatenated Dataset

Given the higher MAE obtained in SFC models compared to concatenated multimodal models (H1.2), we examined in an exploratory analysis whether this difference reflects the information content of the features or merely their number. SFC provides one feature per region, whereas the concatenated multimodal dataset contains the full SC and FC feature sets. We therefore evaluated the MAE of the concatenated datasets as a function of feature set size, separately for each atlas, using the SFC feature count as the baseline.

The results differ substantially by parcellation scheme used. For both the HCP MMP1 and Brainnetome atlas, the concatenated models outperformed the SFC baseline already at equal feature size (Figure 6A, 6C). However, using Gordon’s parcellation framework, the performance of the concatenated dataset was inferior to SFC at equal feature size, but outperformed SFC from twice the SFC feature size onwards (Figure 6B).

**Figure 6.**
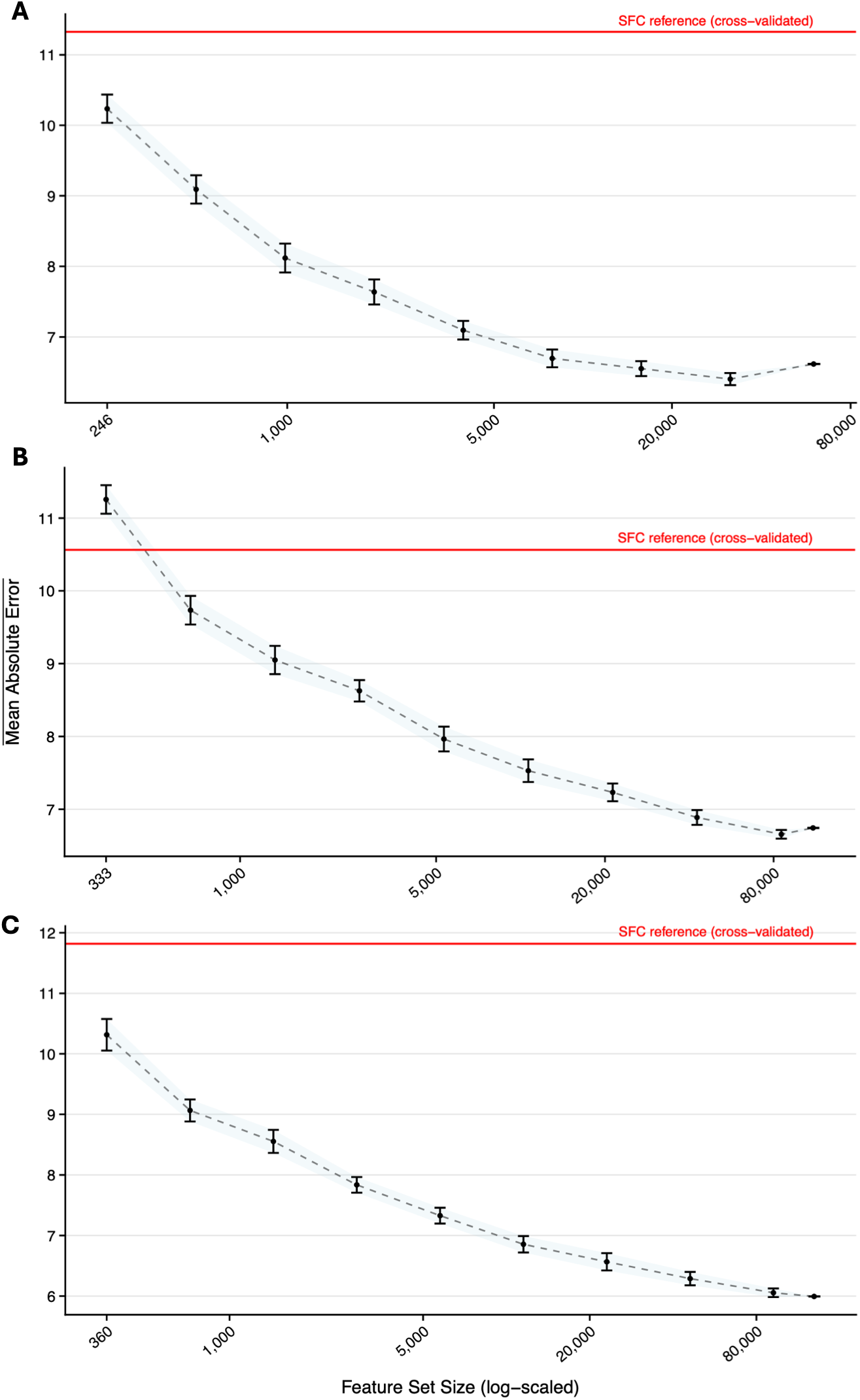
Performance of Concatenated Dataset at Increasing Feature Size. *Note:* y-axis = mean of the mean absolute error; errorbars represent 99% confidence interval; fifty repetitions per feature size with random feature draws were performed; A = Brainnetome, B = Gordon, C = HCP MMP1.

#### Effects of Adjustment Type (RQ2)

The following results are based on the two-way interaction between adjustment and modality. Reported effects reflect conditional main effects, averaged across atlases (see Figure 7). A monotonic decrease in MAE with increasing number of bias adjustment methods is not apparent (H2.1). Comparing no adjustment with one adjustment step for each modality did not reveal significant differences in MAE (all ps > .374). On a similar note, MAE did not differ significantly between two-step or one-step adjustments (all ps > .374). Adding a third adjustment step did not significantly decrease MAE over two step adjustments across different data modalities (all ps > .374).

**Figure 7.**
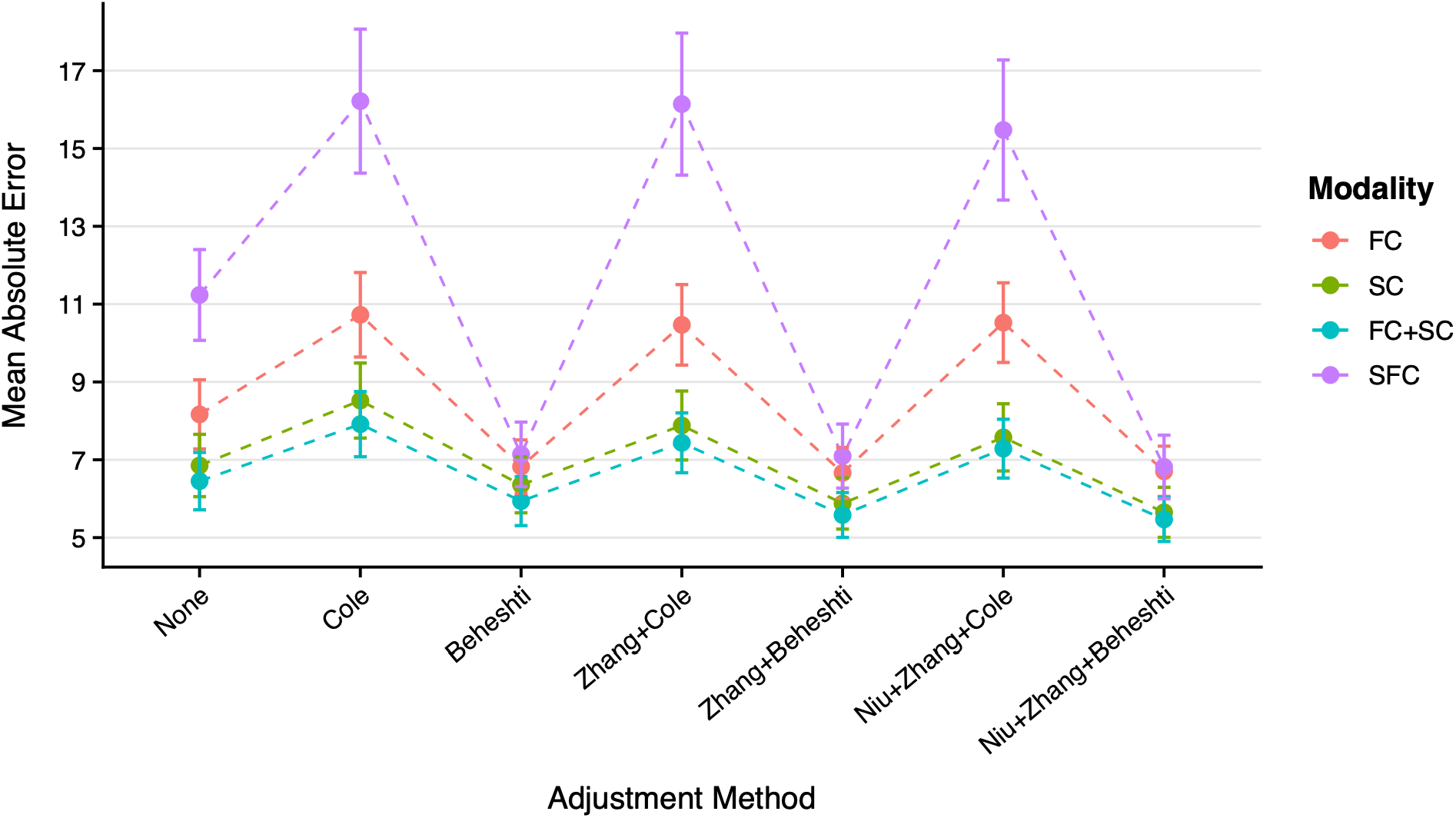
Interaction Plot: Mean Absolute Error by Adjustment x Modality. *Note:* 99% confidence interval used, calculated from raw absolute error for readability (not model-estimated, nor transformed data); FC = functional connectivity, SC = structural connectivity, FC+SC = concatenated structural and functional connectivity, SFC = structure-function coupling.

Using Beheshti sample-level adjustments (i.e., Beheshti only, Zhang + Beheshti, Niu + Zhang + Beheshti) led to significantly lower MAE than using Cole sample level adjustments (i.e., Cole only, Zhang + Cole, Niu + Zhang + Cole) (ps < .001) (H2.2).

#### Effects of Atlas (RQ3)

We examined the simple effects of Atlas within each modality, based on the Modality x Atlas interaction. Reported effects are conditional main effects, averaged across adjustment method (see Figure 8). Since H3 specifies the granularity of the respective atlas (Brainnetome < Gordon < HCP MMP1), not all pairwise but only ordered comparisons are performed. For SC, no significant difference in MAE between the Brainnetome and Gordon nor the Gordon and HCP MMP1 atlas became apparent (all ps > .387). The same was true for the concatenated multimodal models (all ps = .974). For SFC, the Brainnetome atlas led to significantly higher MAE scores than the Gordon atlas (p < .001). However, estimates from the HCP MMP1 and Gordon atlas did not differ significantly (p = .028). For FC, the Gordon atlas yielded significantly higher MAE than both the Brainnetome (p < .001) and HCP MMP1 atlas (p < .01).

**Figure 8.**
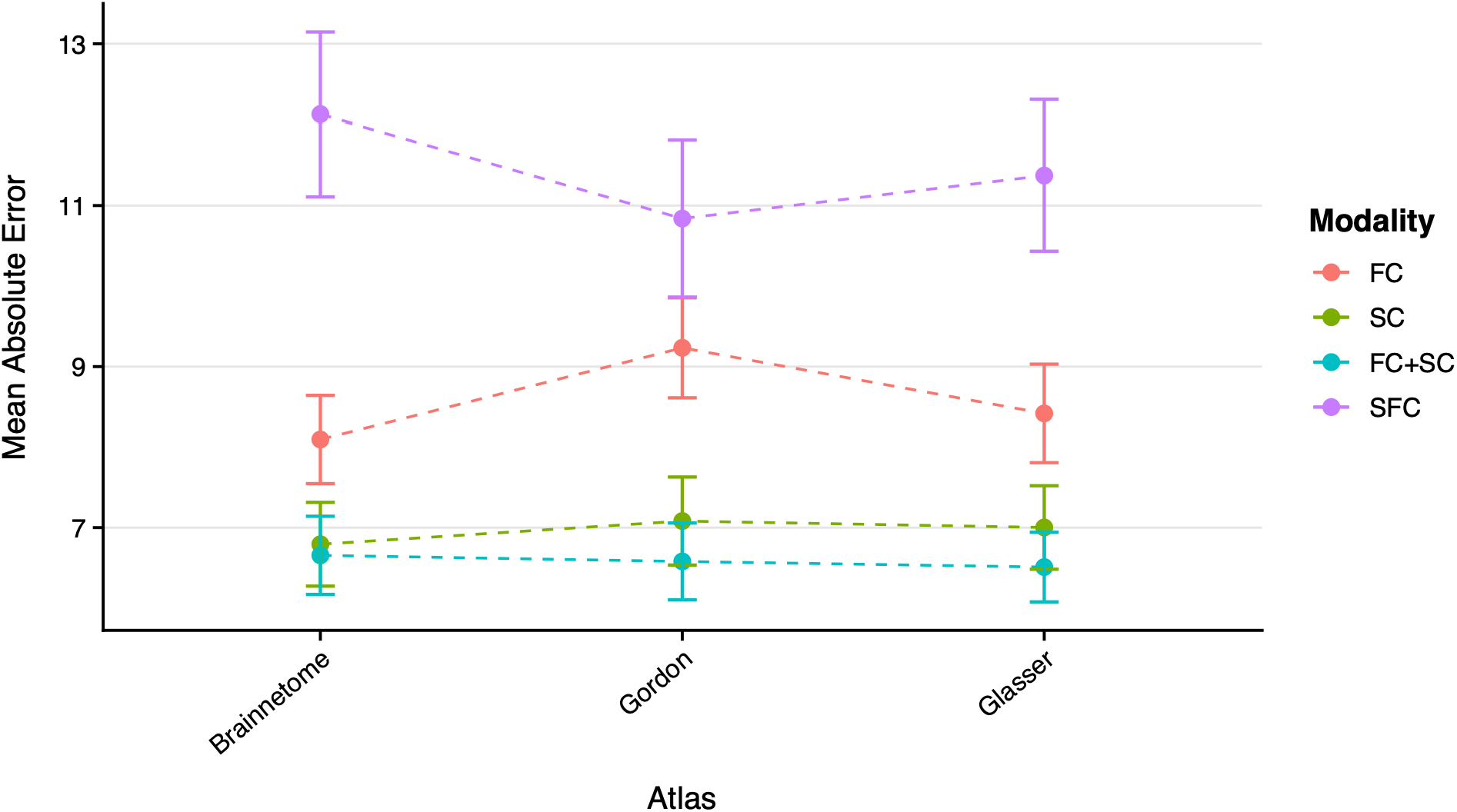
Interaction Plot: Mean Absolute Error by Atlas x Modality. *Note:* 99% confidence interval used, calculated from raw absolute error for readability (not model-estimated, nor transformed data); FC = functional connectivity, SC = structural connectivity, FC+SC = concatenated structural and functional connectivity, SFC = structure-function coupling.

### Regional Overlap in Predictive Nodes (RQ4)

We assigned the connections from the original parcellations (i.e., Gordon, HCP MMP1, and Brainnetome) to the Yeo-7 networks and extracted the top 10 most predictive features. Figure 9 shows circular plots for each atlas × modality combination. High-resolution versions of both the original and reattributed plots are provided in SM.G. Visual inspection suggests that the overall structure of these plots and the degree of communality across atlases within each modality differs, which we quantified with fold enrichment (FE).

**Figure 9.**
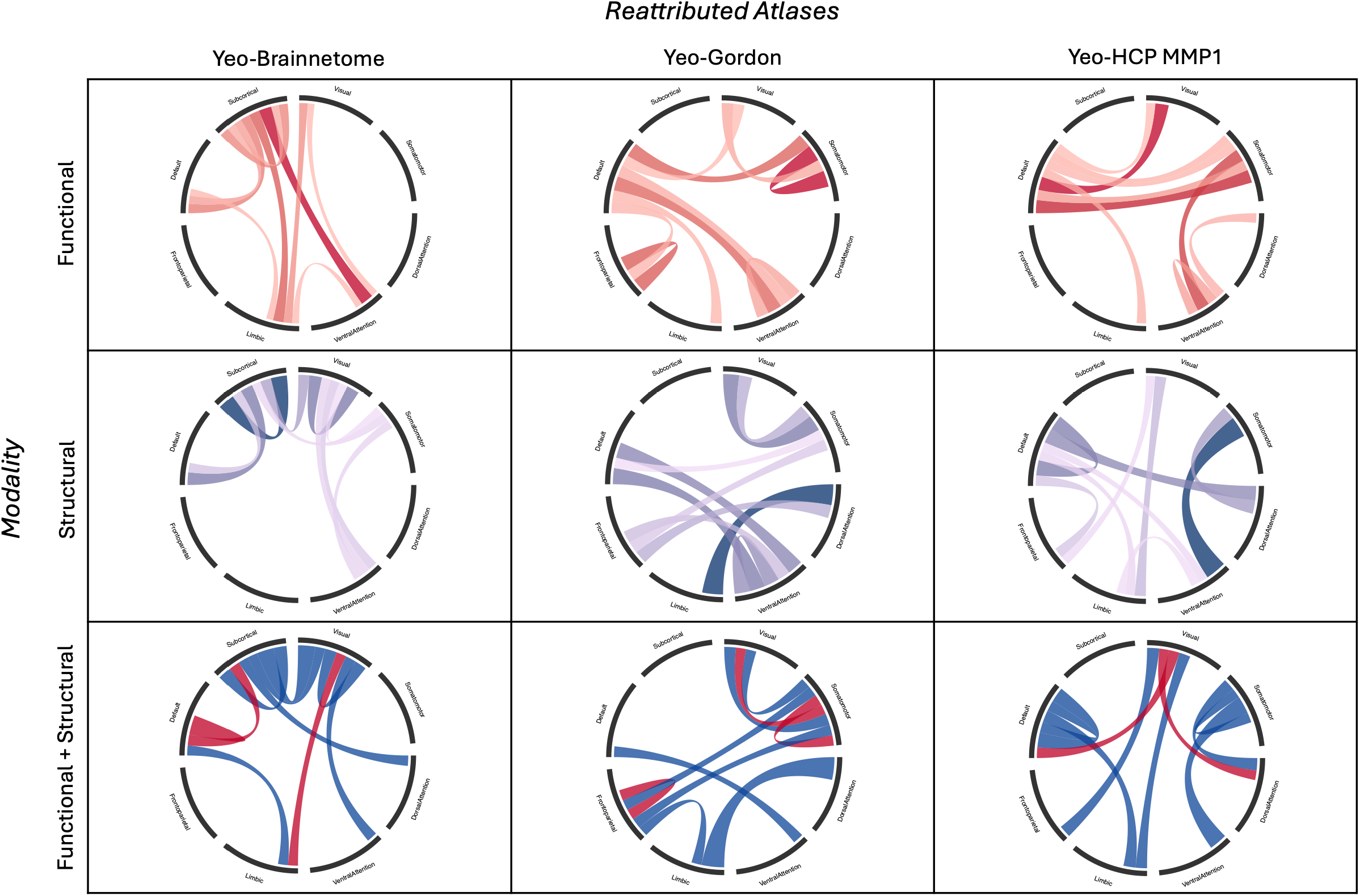
Circular Connectivity Plots for Top 10 Reattributed (Yeo) Predictive Nodes per Atlas. *Notes:* blue = structural connectivity, red = functional connectivity; unimodal models use relative shading by importance from highest to lowest connection within the Top 10;

For SC and the concatenated multimodal model, FE was below 1 across all pairwise and three-way atlas comparisons (overlap below chance level expectation; range 0 – 0.9, see SM.H). Only FC showed FEs exceeding one as outlined in Figure 10. Specifically, the three-way overlap consisted of one shared edge (Default–Limbic; expected overlap = 0.33, FE = 3.0), which was not significant (p = .297). Notably, overlap between the Gordon and HCP MMP1 atlas reached FE of 2.7 using FC features (Expected = 1.5; Overlap = 4; p = .024; regions: Default-Limbic, Default-Somatomotor, Default-Visual, VentralAttention-VentralAttention within top 10 feature edges). Overall, the most predictive connections were largely atlas-specific, in line with H4.

**Figure 10.**
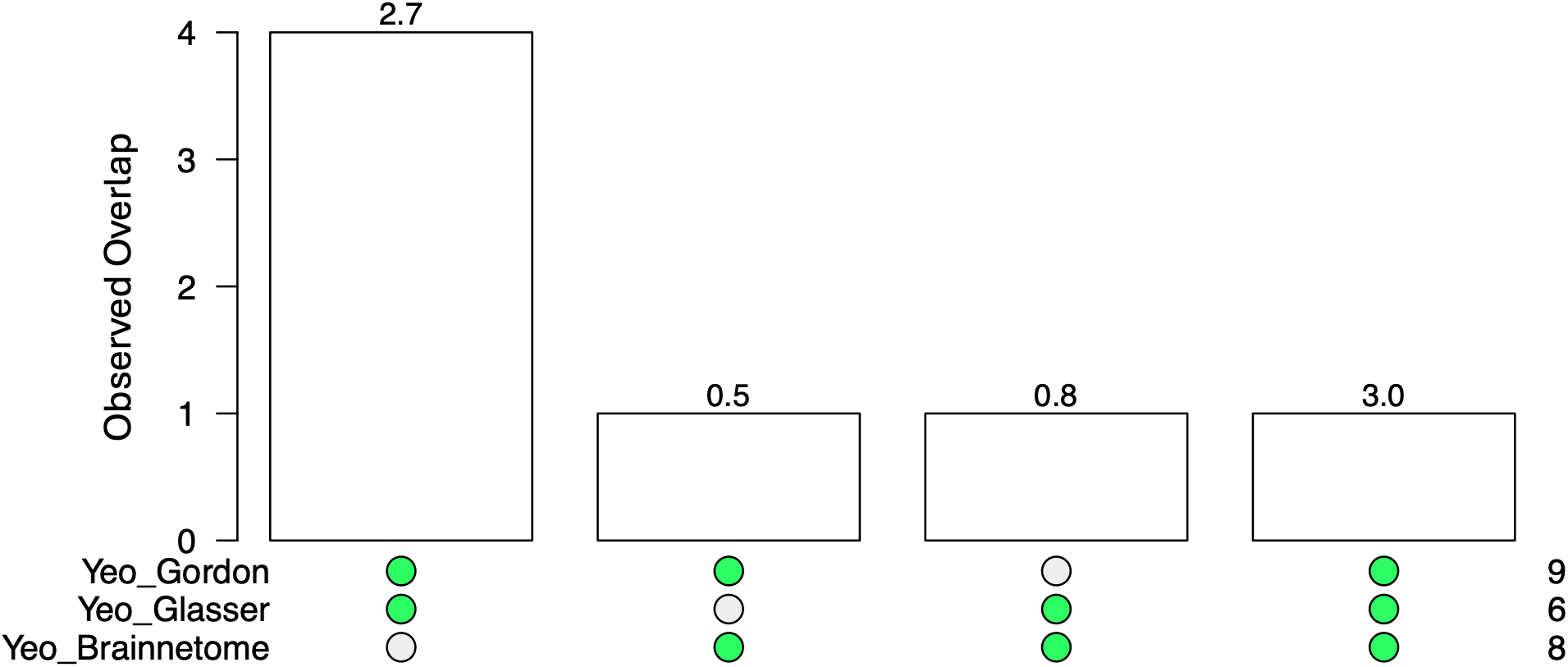
Functional Connectivity: Cross-Atlas Overlap of Top Yeo-7 Edges. *Note:* Bars represent the observed overlap (number of shared network–network edges); numbers above bars indicate fold enrichment (FE = observed / expected overlap under random sampling); green dots denote the atlas combinations included in each intersection; values on the right-bottom indicate the number of unique top-ranked edges per reattributed atlas.

### Exploratory Analysis: Feature Importance

Because the most predictive connections overlapped little across atlases, we examined how feature importance was distributed within each model. We quantified the sparsity of the feature-importance matrices and examined the distribution of non-zero feature importance weights. While matrix sparsity was high across all atlases and modalities (range: 97.78 – 99.43%), many features were non-zero (range 620-833 features). Importance was spread broadly across these features: the top-10 features together captured between 6.03% and 9.57% of the total importance (for age prediction) summed across all non-zero-coefficient features, and 99 to 157 features were needed to reach 50% of the cumulative importance. Since the observed pattern was consistent across atlases and modalities, one representative example is depicted in Figure 13.

**Figure 13.**
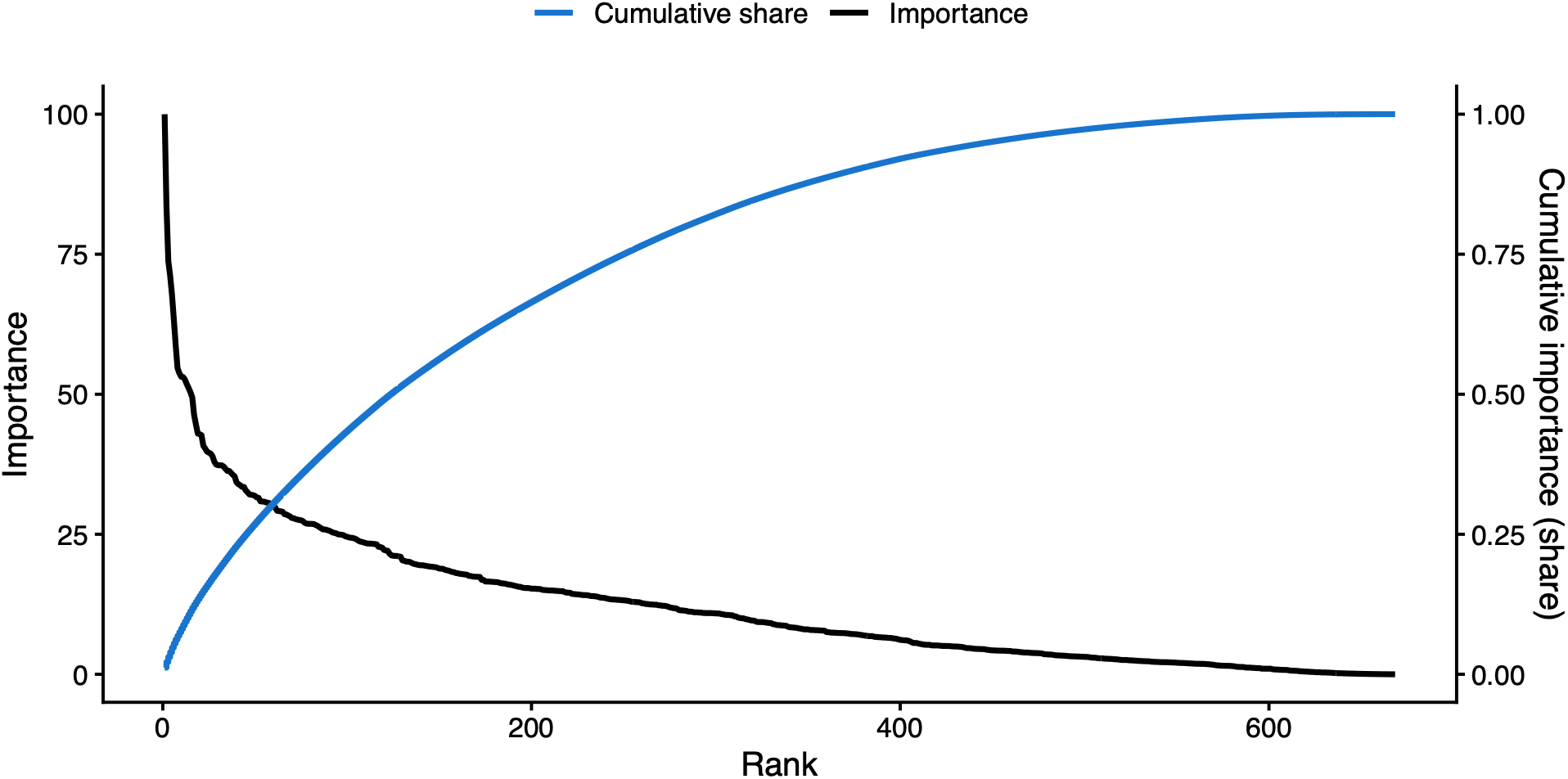
Ranked and Cumulative Feature Importance: Brainnetome Parcellated Functional Connectivity. *Note:* black curve = raw non-zero importance weight per ranked feature (left y-axis), blue curve = cumulative share of total non-zero importance across ranked features (right y-axis); features ranked in descending order of importance weight.

## Discussion

Brain age has emerged as a biomarker of brain health (de Lange et al., 2022) and has been proposed for practical applications, including early detection and prognosis of disorders, evaluation of treatment outcomes, and support of differential diagnosis (Baecker et al., 2021). Its methodological robustness therefore needs to be assessed carefully. In this study, we used a multiverse approach to evaluate the systematic impact of modality (RQ1), bias adjustment technique (RQ2) and atlas choice (RQ3) on the predictive performance (MAE) of brain age estimations. In addition, we examined whether the network-to-network connections contributing most to brain age prediction overlap across atlases in healthy individuals (RQ4).

Across the 84 pipeline conditions, substantial variation emerged in the MAE ranging from 5.26 to 18.30 years and explained variance spanning 44.97% to 90.90%. Modality and adjustment method had medium-sized effects on MAE, and their interaction was of greatest practical relevance. Atlas choice showed no main effect and a significant but negligible interaction with modality. Concatenated multimodal and SC-only models performed best whereas SFC performed worst. Beheshti-based adjustment outperformed Cole-based adjustment and the most predictive connections overlapped little across atlases. Most conditions varied between an MAE of 5 through 8 years, and the wide range was largely driven by a limited subset of poorly performing pipelines. The brain age prediction pipelines examined here are competitive with published benchmarks reported for healthy samples (MAEs of 4.3–13.5 years and correlations of r = 0.43–0.97; Franke & Gaser, 2019). We now discuss each research question and its hypotheses in turn.

Regarding the first research question (RQ1), which examined whether integrating structural and functional information improves brain age prediction, no support was found for either hypothesis. Multimodal integration did not improve prediction relative to unimodal models (H1.1) and SFC performed worse than the concatenated multimodal model rather than better (H1.2). The concatenated multimodal models did not differ significantly from SC alone across adjustment and atlas conditions, and outperformed FC alone in most. SFC led to the highest MAE in most conditions. These findings suggest that SC captured most of the information relevant for predicting chronological age, and that FC added little beyond it. One possible explanation is that the age-related information contained in FC overlaps with that captured by SC (cf., Damoiseaux, 2017). Another is that FC may contain a weaker age-related signal than SC, limiting its incremental predictive value. Although the specific reasons why FC provides limited additional predictive information has not been directly investigated, previous brain-age studies have consistently found weaker age prediction from FC than from structural measures (de Lange et al., 2020; Jirsaraie, Gorelik, et al., 2023). The inferior performance of SFC was not a matter of feature size alone, since at equal feature sizes the concatenated models still outperformed SFC across two out of three parcellation frameworks. How structural and functional information is represented therefore appears to matter. Computing structure-function coupling may discard predictive information or introduce additional variability, and regional coupling may in itself not be a strong or stable marker of chronological age. The present findings thus do not rule out that SC and FC carry complementary information, but they show that neither simple concatenation nor the SFC representation used here translated into better prediction of chronological age compared to SC alone. Multimodal benefit could therefore be context-dependent and contingent on how features are combined, rather than an inherent property of larger input spaces. This echoes Jirsaraie, Gorelik, et al. (2023) who reported that multimodal models tend to be more accurate when it comes to chronic brain disorders, whereas unimodal functional models can be sensitive to a variety of phenotypes. Together, these observations caution against the default adoption of multimodal pipelines and call for more targeted empirical validation of when and why combining modalities improves performance.

Turning to bias-adjustment approaches (RQ2), increasing the number of correction steps did not produce a significant monotonic improvement in predictive accuracy (H2.1), indicating that more adjustment is not synonymous with better prediction. This finding is consistent with the primary purpose of bias adjustment. These procedures are designed to reduce systematic age-related bias estimates rather than to directly minimize prediction error (Zhang et al., 2023). Consequently, reducing bias does not necessarily translate into improved prediction accuracy. The type of adjustment did matter: Beheshti-based corrections consistently outperformed Cole-based ones across all conditions (H2.2), in line with Zhang et al. (2023). A likely reason is that Cole’s rescaling of the predictions amplifies the noise alongside the signal, whereas the Beheshti approach does not. For researchers whose primary goal is maximizing prediction accuracy in healthy samples, Beheshti-based adjustments therefore appear to be the most suitable choice.

The third factor, atlas choice (RQ3), had little influence on accuracy. The hypothesis that finer parcellations yield lower error (H3) was not supported. Atlas choice has been of much concern in the literature (cf. Bryce et al., 2021), but its effect on MAE was, although significant, negligible. The atlas differences that emerged did not follow the granularity hypothesis. No clear pattern linked more granular atlases to lower MAE. Atlas choice thus appears to matter little for predictive accuracy in this setting, although it does not settle whether it impacts conclusions about which brain regions, areas or networks drive prediction.

The question whether the most predictive connections are consistent across atlases was addressed by RQ4, which asked whether the most predictive connections remain consistent across atlases. The hypothesis (H4) that connections vary across atlases was supported. When the three atlases (i.e., Brainnetome, Gordon and HCP MMP1) were embedded in broader Yeo7 networks, the overlap between the 10 most predictive features was limited, irrespective of whether structural and functional connectivity were considered separately or jointly. Moreover, evidence arose that instead of the top 10 connections, rather approximately 500 unique connections are relevant for accurate brain age predictions. The low overlap may therefore reflect arbitrary selection among similarly weighted features as much as genuine atlas dependence. It is consistent with brain aging reflecting a distributed, multifactorial neurobiological change rather than a process confined to discrete regions or networks (Raz & Rodrigue, 2006).

Given these results, researchers might be tempted to conclude that brain age prediction in healthy participants should (a) use concatenated structural and functional data, (b) apply Beheshti-based adjustment methods, and (c) select a parcellation framework based on convenience or theoretical preference. While defensible on purely predictive grounds, this recommendation falls short of addressing the ongoing tension between prediction accuracy and biomarker utility (Jirsaraie, Gorelik, et al., 2023; Jirsaraie, Kaufmann, et al., 2023; Schulz et al., 2025). With the best-performing pipeline capturing over 90% of the variance in chronological age, relatively little residual variance remains. Yet this residual is precisely what constitutes the brain age gap (BAG); the quantity on which clinical applications depend. Within this narrow band of unexplained variance, genuine biological signal reflecting individual differences in aging trajectories (due to brain aging) must be separated from methodological noise introduced by model imprecision, preprocessing choices, and measurement error. This has been defined as the biomarker paradox (Salih et al., 2023): as predictive accuracy improves, the brain age gap shrinks, and distinguishing meaningful individual variation from artifact might become increasingly difficult. In other words, reducing prediction error to zero would eliminate the brain age gap, rendering the biomarker meaningless since it would simply represent chronological age. Critically, Schulz and colleagues (2025) complicated the picture further beyond smaller residual variance. Their analysis of over 46,000 UK Biobank participants demonstrated that optimizing models for chronological age prediction actively diminished their sensitivity to disease-relevant variation. Specifically, advanced accuracy-optimized models (i.e., neural networks) tended to rely on stable, low-variance features that track tightly with age but carry little information about pathology. Thus, according to the authors and as utilized in this study, regularized regressions might be more beneficial (Schulz et al., 2025). Consequently, predictive performance does not necessarily equate to clinical relevance (Jirsaraie, Kaufmann, et al., 2023). Following these thoughts on the biomarker paradox, a major obstacle in brain age predictions is the absence of a ground truth.

Nonetheless, maximizing prediction accuracy should not be dismissed entirely. Firstly, large individual prediction errors make single-subject inference unreliable. Individual errors can range from near-perfect prediction to discrepancies exceeding 25 years (Lancaster et al., 2018). Thus, narrowing this gap is essential to reduce noise. Moreover, it has been argued that reaching MAE values of 0 that would deem the biomarker redundant is unrealistic (Lancaster et al., 2018). Secondly, more reliable (low-MAE) brain age predictions have shown to monitor the progression of long-term chronological age in healthy subjects well (Dörfel et al., 2023). Thirdly, under quality constraints (i.e., lower quality scans) more accurate models were of greater use despite lower chance of detecting cognitive differences (Jirsaraie, Kaufmann, et al., 2023). This suggests that accuracy and utility are only partially dissociable performance dimensions that may require semi-independent optimization depending on the research goal.

This investigation contributes to this debate by demonstrating that pipeline decisions (spanning data modality, bias-adjustment method and parcellation) collectively produce meaningful variation in prediction accuracy. This variance is not merely noise: it maps onto a space in which researchers can, in principle, position their pipeline according to their primary research goal. Where the aim is predictive precision in healthy samples, such as establishing normative brain age distributions or quantifying individual deviation from a healthy aging trajectory, the present findings support prioritizing concatenated multimodal data or structural connectivity data and Beheshti-based adjustment. Where the aim is biomarker sensitivity, the accuracy-sensitivity tension documented in the literature suggests that pipelines maximizing predictive fit may not be optimal, though the present study does not test sensitivity to pathological variation. As a consequence, we cannot recommend specific combinations for that purpose since the appropriate amount of accuracy while retaining sensitivity is yet unknown.

Several avenues for future research emerge from the present findings. First, the multiverse examined here could be expanded to include additional researcher degrees of freedom. Prior investigations considered preprocessing pipelines (Dafflon et al., 2022) as well as the selection of machine learning algorithms (More et al., 2023). However, feature selection strategies and sample composition might introduce further variability into brain age estimates. Second, the present findings should be extended to clinical samples. A pipeline that minimizes MAE in healthy individuals does not necessarily produce a BAG that is most sensitive to disease-related deviations in brain aging. Applying the present multiverse approach to clinical cohorts would therefore reveal which pipeline configurations yield a BAG appropriate in predicting and thus capturing disease-related brain changes. Third, longitudinal validation is essential: a model’s ability to track within-person change over time is relevant to clinical applications and the pipeline choices that optimize cross-sectional prediction may not be the same as those that yield the most stable and sensitive longitudinal brain age trajectories. Lastly, establishing ground-truth benchmarks for brain aging (through convergence with neuropathological, genetic, or epigenetic markers) would provide an external criterion against which the biological validity of different pipeline configurations could be assessed.

Despite efforts to minimize bias through systematic design and methodological transparency, several limitations warrant acknowledgment. Only a single machine learning algorithm was employed, which constrains the generalizability of the findings to other methods (particularly non-linear or deep learning approaches that may interact differently with modality and atlas choices). Nevertheless, this algorithm was deliberately selected for its favorable position along the accuracy-utility trade-off, retaining larger effect sizes than more complex models in brain age prediction, as demonstrated by Schulz et al. (2025). Furthermore, this study does not engage with the ongoing debate surrounding the ground truth of brain aging within its study design. This question is of considerable importance for establishing appropriate benchmarks against which pipelines’ predictive performance can be evaluated. In greater detail, it is difficult to judge what level of prediction error constitutes biologically meaningful deviation versus methodological noise. Many questions therefore remain regarding optimal sample constitution, ground truth in brain age estimation, and the extension of multiverse approaches to clinical contexts.

Nonetheless, the present study provides a rigorous methodological investigation of a multitude of seemingly routine decisions within brain age prediction pipelines. The results demonstrate that such decisions carry meaningful consequences for predictive performance and possibly the validity of downstream inferences. As the field moves toward clinical utility, thoughtful pipeline selection guided by research objectives rather than convenience becomes not merely good practice but a methodological necessity.

## Data Availability and Reproducibility Statement

All quarto scripts used for statistical analyses will be made publicly available on GitHub upon journal publication.

## Conflict of Interest Statement

The authors declare no conflict of interest.

## Acknowledgments

Data used in this study were obtained from the Cambridge Centre for Ageing and Neuroscience (CamCAN) repository (available at www.mrc-cbu.cam.ac.uk/datasets/camcan/). CamCAN data collection and sharing was supported by the UK Biotechnology and Biological Sciences Research Council (grant number BB/H008217/1), the UK Medical Research Council, and the University of Cambridge.

## Author Contributions

J.F.: Conceptualization, Methodology, Software, Validation, Visualization, Formal analysis, Writing: Original Draft, Review and Editing

W.D.B.: Conceptualization, Methodology, Supervision, Preprocessing, Writing: Review and Editing

L.S.: Conceptualization, Writing: Review and Editing

## Notes

http://www.mrc-cbu.cam.ac.uk/datasets/camcan/

### Competing Interest Statement

The authors have declared no competing interest.

