## Supplementary Materials for "A Multiverse Approach to Brain Age Prediction: Exploring the Impact of Modality, Bias Adjustment, and Atlas"

### Supplemental Material A (SM.A)

### Preliminary Analysis: Selection of Machine Learning Method

Functional connectivity data from 560 participants, parcellated using the Gordon atlas, were used for a preliminary comparison of machine learning approaches. A regularized regression model (elastic net) was evaluated alongside several alternative methods, including ordinary least squares regression, a shallow neural network, gradient boosting, and both linear and nonlinear support vector machines.

The comparison between elastic net and the competing methods was intentionally asymmetrical. The competing models could not be reliably fitted on the full high-dimensional functional connectivity data due to convergence and computational limitations. As a result, these models were trained on reduced representations obtained using principal component analysis (PCA; see A1). Direct feature selection in the original connectivity space (i.e., without using principal components) was not considered appropriate as each individual connection captures only a very small proportion of the total variance, such that selecting a limited subset of features would explain little meaningful variance overall. To maximize their predictive performance, the top 50 principal components most strongly associated with age were selected (distribution tails in A2). This procedure is expected to advantage PCA-based models relative to elastic net, which was trained directly on the original feature space and thus selected relevant features itself (regularization).

Predictive accuracy was evaluated using mean absolute error (Table A1). Under these conditions, elastic net achieved predictive performance comparable to that of PCA-based models. Given the overall similarity in performance and the fact that interpretability of individual functional connections is retained only in elastic net, because it operates on the original connectivity features, regularized regression was selected for all subsequent analyses.

Moreover, simpler machine learning models incorporating regularization (such as elastic net) have been shown to yield more meaningful brain age biomarkers and to retain larger effect sizes than more complex machine learning approaches (Schulz et al., 2025).

**Table A1**

*Preliminary Comparison of Machine Learning Approaches for Brain Age Prediction*

| Model | MAE | $R^2$ | RMSE |
|---|---|---|---|
| Non-Linear Support Vector Machine (PCA) | 8.32 | 0.70 | 10.40 |
| Linear Regression (PCA) | 8.63 | 0.70 | 10.71 |
| Elastic Net (*original feature set*) | 8.65 | 0.68 | 10.84 |
| Linear Support Vector Machine (PCA) | 8.73 | 0.69 | 10.87 |
| Gradient Boosting (PCA) | 9.27 | 0.63 | 11.79 |
| Shallow Neural Network (PCA) | 12.16 | 0.50 | 15.74 |

*Note:* RMSE = root mean square error, MAE = mean absolute error; dataset = functional connectivity data of 560 participants; (PCA) = fit on 50 highest correlated principal components to age; 5-fold cross-validation has been applied to all methods.

**Figure A1**

*Cumulative Variance Graph of Principal Components based on Train Set from Functional Connectivity Data (Gordon-parcellations)*

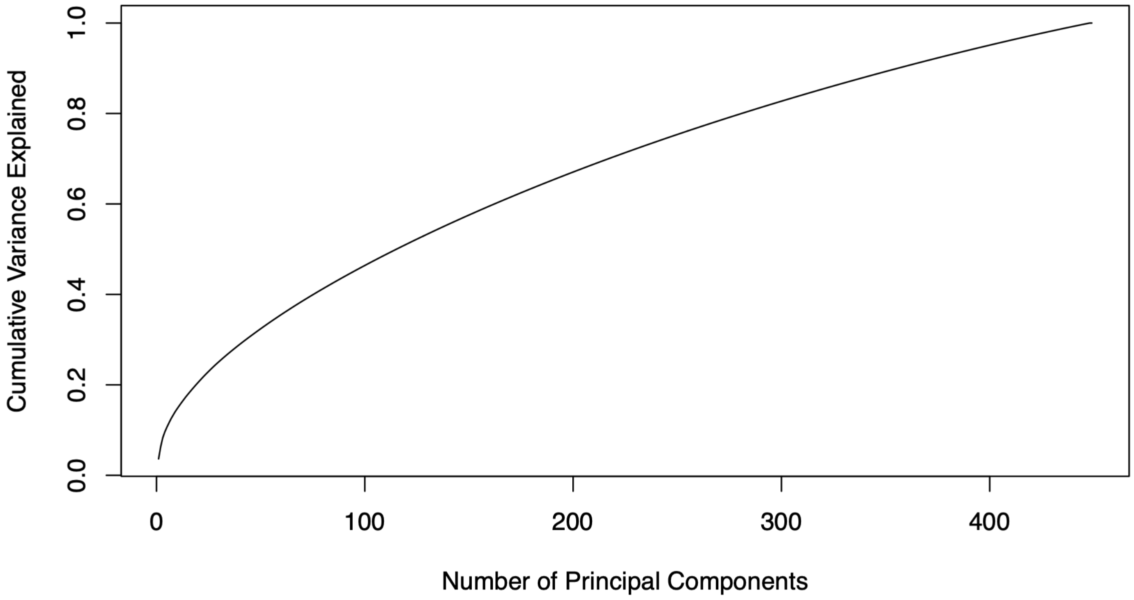

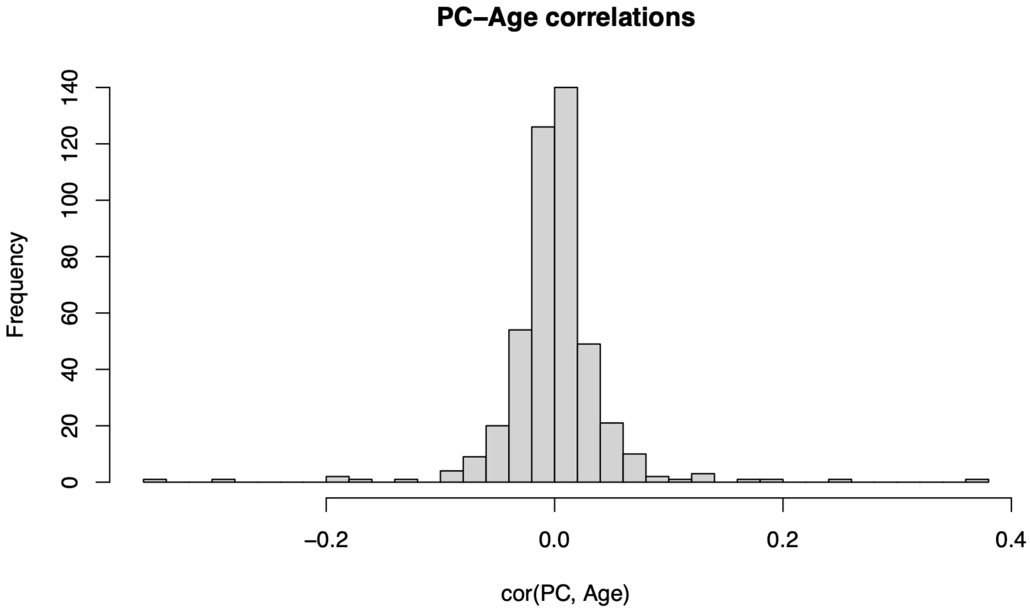
PC−Age correlations
Frequency
0
20
40
60
80
100
120
140
−0.2
0.0
0.2
0.4
cor(PC, Age)

### Supplemental Material B (SM.B)

### Elastic Net Regression

Elastic net estimates were conceptualized in the following manner (cf. Friedman et al., 2010):

$$\min_{\beta_0, \beta} \left\{ \frac{1}{2N} \sum_{i=1}^{N} (y_i - \beta_0 - x_i^T * \beta)^2 + \lambda \left[ \frac{1-\alpha}{2} * \|\beta\|_{\ell 2}^2 + \alpha * \|\beta\|_{\ell 1} \right] \right\}$$

where $y_i$ denotes chronological age of subject i,

$x_i \in \mathbb{R}^p$ represents the vector of p neuroimaging features,

$\beta \in \mathbb{R}^p$ represents corresponding coefficient vector,

$\lambda \geq 0$ represents the overall regularization strength and

$\alpha \in [0,1]$ the ridge-lasso mixing parameter.

### Supplemental Material C (SM.C)

### Bias-Adjustment Methods

Procedure

Each adjustment procedure is described in detail below, including its conceptual rationale and mathematical formulation. Equations are provided in a generic-single observation style. Thus, the index for observations (i) was omitted to sustain readability. For the Zhang adjustment observation indexing has been included since it is relevant for the adjustment's conceptualization. The predicted brain age gap (PAD) and brain age gap (BAG) refer to the same concept; since the bias adjustment literature favours the former term, it was used throughout.

The sample level bias adjustment originally proposed by Cole first regresses the predicted age ($\hat{Y}$) on the observed age (Y) to capture linear bias. Following this the bias is reduced by removing the intercept ($\beta_{pred}$) from the age-predictions and dividing by the slope ($\alpha_{pred}$). As a result, the adjusted predicted age gap ($PAD_c$) is obtained (Zhang et al., 2023). Lastly, the adjusted predicted age ($\widehat{Y_c}$) is created by adding the adjusted predicted brain age ($PAD_c$) to the chronological age (Y).

$$\hat{Y} = \alpha_{pred} * Y + \beta_{pred} \quad \text{(C1)}$$

$$PAD_c = \widehat{Y_c} - Y = \frac{\hat{Y} - \beta_{pred}}{\alpha_{pred}} - Y \quad \text{(C2)}$$

$$\widehat{Y_c} = PAD_c + Y \quad \text{(C3)}$$

In Beheshti's method of sample level bias adjustment, the predicted age difference (PAD) is modelled as a linear function of observed age ($Y$). Following this, the fitted age gap trend is subtracted from the predicted age ($\hat{Y}$). Lastly, the adjusted predicted age ($\widehat{Y_c}$) is extracted. Consequently, Cole and Beheshti adjustment are similar, only that Cole's fits a regression on the predicted age, inevitably containing higher variance usually leading to lower MAE predictions (Zhang et al., 2023).

$$PAD = \alpha_{pad} * Y + \beta_{pad} \quad \text{(C4)}$$

$$PAD_c = \widehat{Y_c} - Y = \hat{Y} - ([\alpha_{pad} + 1] * Y + \beta_{pad}) \quad \text{(C5)}$$

$$\widehat{Y_c} = PAD_c + Y \quad \text{(C6)}$$

The proposed age-level specific bias adjustment aims at removing residual bias within each exact age cohort. This is achieved by subtracting the average corrected PAD ($\mu_a$) for each age bin ($a$) from the individual ($i$). This creates age-conditional corrected predicted age gap ($PAD_i^{ac}$). This can then be used to adjust the predicted age of each individual ($Y_c$) (Zhang et al., 2023). However, in the present study, age cohorts were defined as three-year age ranges rather than single-year values. The bin width was selected after examining the age distribution in the training and test data to ensure sufficient sample size within each cohort while preserving age specificity.

$$PAD_i^{ac} = PAD_i - \mu_a \qquad \text{(C7)}$$

$$\widehat{Y_c} = PAD_i^{ac} + Y \qquad \text{(C8)}$$

Furthermore, Niu et al. (2019) proposed that age-dependent bias in brain-age prediction may follow nonlinear patterns and therefore may not be fully removed by linear correction procedures alone. To capture higher-order bias structure, the pre-corrected brain-age gap ($PAD$) was modeled as a polynomial function of chronological age (including a quadratic term) and its interactions with sex ($\widehat{PAD}$). The nonlinear-adjusted gap was defined as the residual of this model, i.e., the difference between the observed gap and the fitted nonlinear bias component ($PAD_c = \varepsilon_{\widehat{PAD}}$). Finally, corrected predicted age was reconstructed by adding chronological age (Y) to the adjusted gap ($PAD_c$). Note: Within this study only the quadric effect of age, the linear effect of sex and the interaction between sex and quadratic age has been removed to avoid reintroducing linear age bias / over-adjusting for linear age bias.

$$\widehat{PAD} = \beta_0 + \beta_1 * Y + \beta_2 * Y^2 + \beta_3 * sex + \beta_4 * sex * Y + \beta_5 * sex * Y^2 \qquad \text{(C9)}$$

$$PAD_c = PAD - \widehat{PAD} \qquad \text{(C10)}$$

$$\widehat{Y_c} = PAD_c + Y \qquad \text{(C11)}$$

<u>Implications of Bias Adjustment</u>

It is important to note that chronological age entered the prediction pipeline at two distinct stages. First, it was used to compute the brain age gap itself (predicted age − chronological age), while the underlying model prediction remained fully label-free. Second, bias-adjustment procedures used chronological age to model and remove systematic age-dependent trends in the gap. Consequently, MAE evaluated on the corrected brain age gap reflects residual error after age-dependent calibration rather than label-free prediction accuracy. This does not undermine the purpose of this comparison since the goal is not to isolate raw label-free prediction accuracy, but to assess how different bias-adjustment strategies affect the brain age gap (by MAE).

### Supplemental Material D (SM.D)

### Multilevel Model

$$Sqrt(AE_{ij}) = \beta_{0j} + \beta_{1j} * Modality_{ij} + \beta_{2j} * Adjustment_{ij} + \beta_{3j} * Atlas_{ij} + \beta_{4j} * Modality_{ij} * Adjustment_{ij} + \beta_{5j} * Adjustment_{ij} * Atlas_{ij} + \beta_{6j} * Modality_{ij} * Atlas_{ij} + \beta_{7j} * Modality_{ij} * Adjustment_{ij} * Atlas_{ij} + \varepsilon_{ij} \quad \text{(D1)}$$

$$\varepsilon_{ij} \sim N(0, \sigma_{\varepsilon}^2) \quad \text{(D2)}$$

$$\beta_{0j} = \gamma_{00} + \upsilon_{0j} \quad \text{(D3)}$$

$$\upsilon_{0j} \sim N(0, \sigma_{\upsilon_0}^2) \quad \text{(D4)}$$

$$\beta_{1j} = \gamma_{10} \quad \text{(D5)}$$

$$\beta_{2j} = \gamma_{20} \quad \text{(D6)}$$

$$\beta_{3j} = \gamma_{30} \quad \text{(D7)}$$

$$\beta_{4j} = \gamma_{40} \quad \text{(D8)}$$

$$\beta_{5j} = \gamma_{50} \quad \text{(D9)}$$

$$\beta_{6j} = \gamma_{60} \quad \text{(D10)}$$

$$\beta_{7j} = \gamma_{70} \quad \text{(D11)}$$

$$Sqrt(AE_{ij}) = \gamma_{00} + \gamma_{10} * Modality_{ij} + \gamma_{20} * Adjustment_{ij} + \gamma_{30} * Atlas_{ij} + \gamma_{40} * Modality_{ij} * Adjustment_{ij} + \gamma_{50} * Adjustment_{ij} * Atlas_{ij} + \gamma_{60} * Modality_{ij} * Atlas_{ij} + \gamma_{70} * Modality_{ij} * Adjustment_{ij} * Atlas_{ij} + \upsilon_{0j} + \varepsilon_{ij} \quad \text{(D12)}$$

### Supplemental Material E (SM.E)

### Power Analysis

Conventional software for power and sensitivity analysis, such as G*Power (Faul et al., 2007), assumes independence of observations. This assumption is violated in multilevel models when the intraclass correlation coefficient (ICC) is nonzero. Nevertheless, the multilevel model chosen for this study includes a random intercept only and is therefore relatively simple. For such models, power can be approximated by applying a corrected, or "effective" sample size (Hox et al., 2010).

Accordingly, the power analysis followed a three-step approximation for simple multilevel designs:
(1) estimation of the design effect based on the ICC,
(2) computation of an effective sample size, and
(3) power analysis using standard power analysis software.

(1) Intraclass Correlation Coefficient (ICC).
Reported ICC estimates for functional and structural connectivity data vary substantially, ranging from approximately 0.29 (Noble et al., 2019) to 0.75 (Tsai, 2018). Because the planned multilevel analyses include structural, functional, and coupled connectivity measures, power was evaluated under both the lowest reported ICC (best-case scenario) and highest reported ICC (worst-case scenario) to reflect plausible bounds of clustering.

(2) Effective Sample Size
The effective sample size has been determined as specified in the handbook "Multilevel Analysis: Techniques and Applications" (Hox et al., 2010, p. 5) for both ICC scenarios.

In which

- $n$ denotes the total number of observations
- $n_{cluster}$ denotes the cluster size
- $\rho$ denotes the intraclass correlation

Applying the variation of the Kish formula (Kish, 1965) outlined above, leads to following effective sample sizes:

(3) Power Analysis in G*Power.

Power analyses were conducted using G*Power (Faul et al., 2007) by specifying fixed-effects ANOVA models corresponding to each factor included in the multilevel model. Because the dependency structure was already incorporated through the effective sample size adjustment, a fixed-effects framework was used rather than a repeated-measures ANOVA. Power was evaluated across small (f = 0.10), medium (f = 0.25), and large (f = 0.40) effect sizes under both ICC scenarios. Power Analysis has only been conducted for the main effects (not interaction terms) since these encapsulate the primary interest of this study.

Table E1

*Power Estimation given Effect Size for Factor Modality (4 levels)*

| Expected Intraclass Correlation | Expected Effect Size | | |
|---|---|---|---|
| | f = .1 | f = .25 | f = .4 |
| ICC = .29 | .15 | .95 | 1 |
| ICC = .75 | .05 | .47 | .95 |

Table E2

*Power Estimation given Effect Size for Factor Adjustment Type (7 levels)*

| Expected Intraclass Correlation | Expected Effect Size | | |
|---|---|---|---|
| | f = .1 | f = .25 | f = .4 |
| ICC = .29 | .09 | .89 | 1 |
| ICC = .75 | .03 | .33 | .88 |

Table E3

*Power Estimation given Effect Size for Factor Atlas (3 levels)*

| Expected Intraclass Correlation | Expected Effect Size | | |
| --- | --- | --- | --- |
| | f = .1 | f = .25 | f = .4 |
| ICC = .29 | .18 | .97 | 1 |
| ICC = .75 | .06 | .55 | .97 |

### Supplemental Material F (SM.F)

### Explained Variance Plots

Figure F1

*Heat-Map of Explained Variance (R2) Across 84 Experimental Conditions*

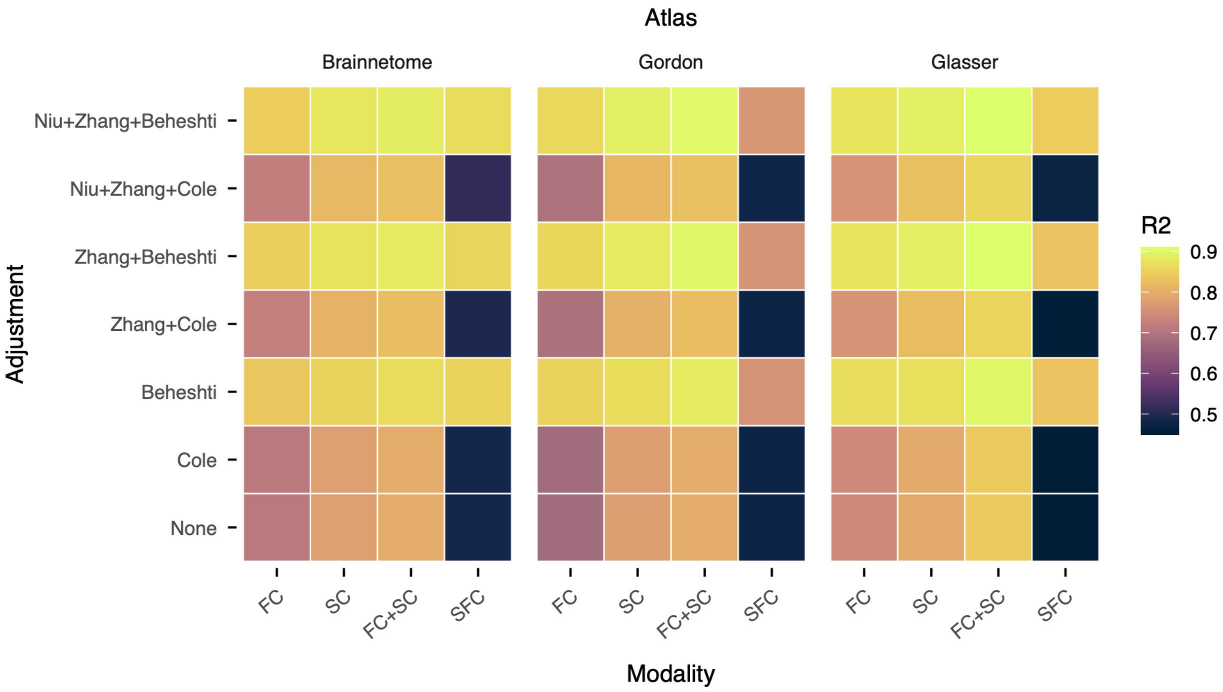


*Note:* FC = functional connectivity, SC = structural connectivity, FC+SC = concatenated structural and functional connectivity, SFC = structure-function coupling.

Figure F2

*Frequency Distribution of Explained Variance (R2) Across Conditions*

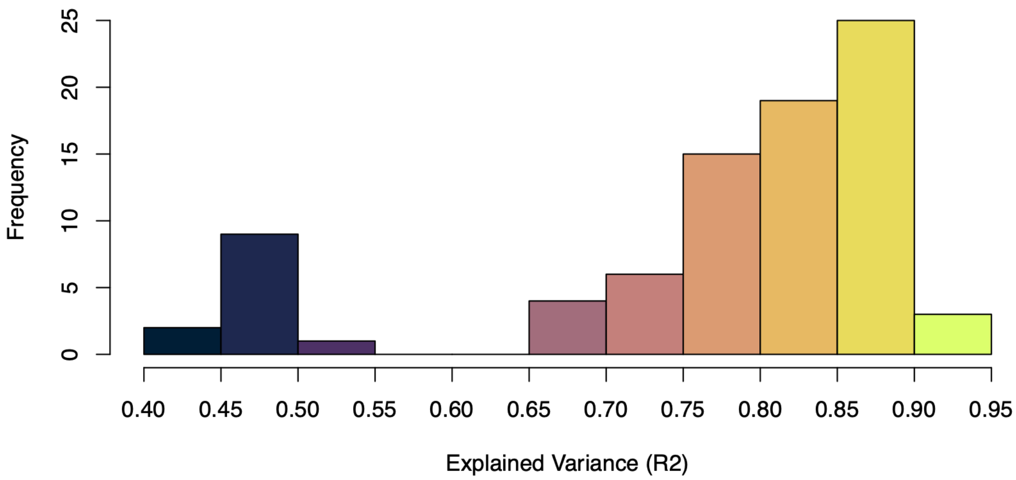

### Supplemental Material G (SM.G)

### Circular Plots of Feature Importance (Brain Regions)

Figure G1

*Top 10 Reattributed (Yeo) Predictive Nodes: Functional Connectivity, Brainnetome Atlas*

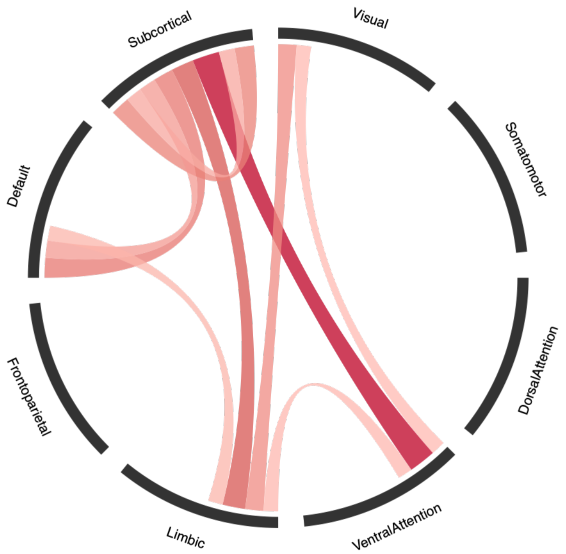


Figure G2

*Top 10 Reattributed (Yeo) Predictive Nodes: Structural Connectivity, Brainnetome Atlas*

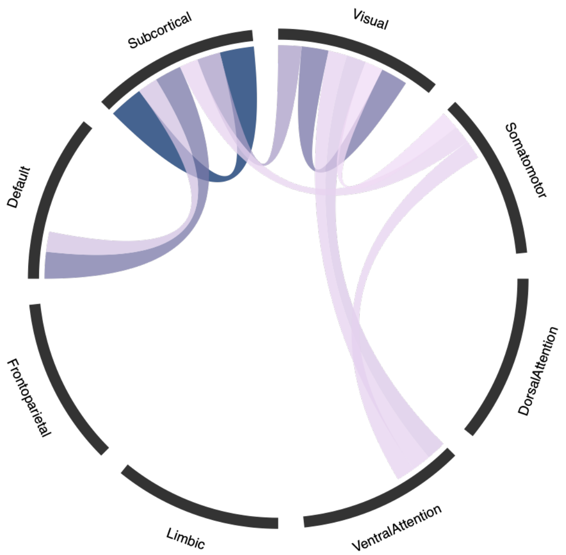

Figure G3

*Top 10 Reattributed (Yeo) Predictive Nodes: Functional Connectivity, HCP MMP1 Atlas*

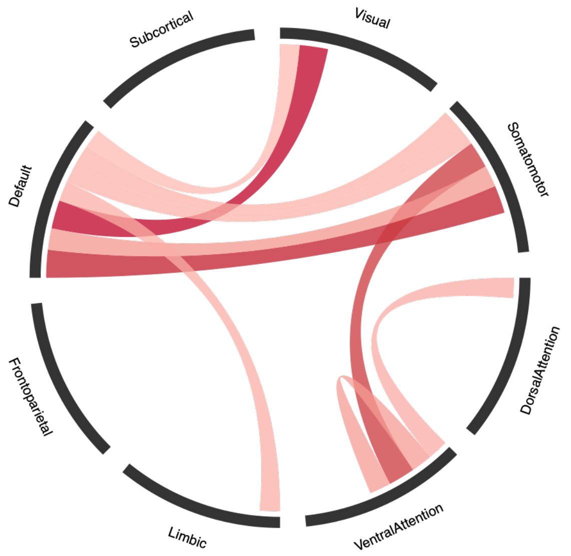


Figure G4

*Top 10 Reattributed (Yeo) Predictive Nodes: Structural Connectivity, HCP MMP1 Atlas*

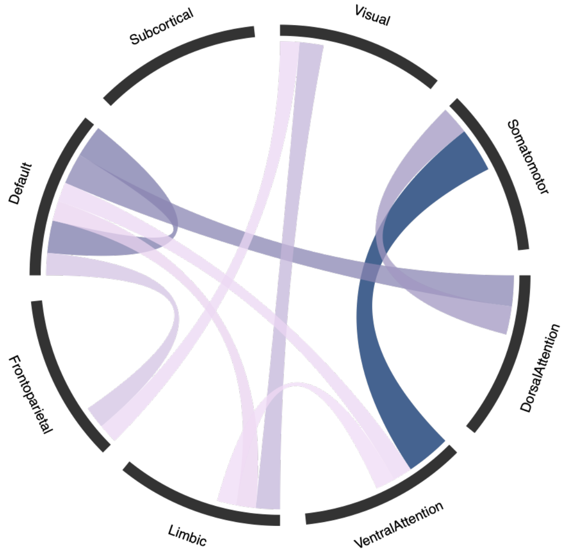


Figure G5

*Top 10 Reattributed (Yeo) Predictive Nodes: Functional Connectivity, Gordon Atlas*

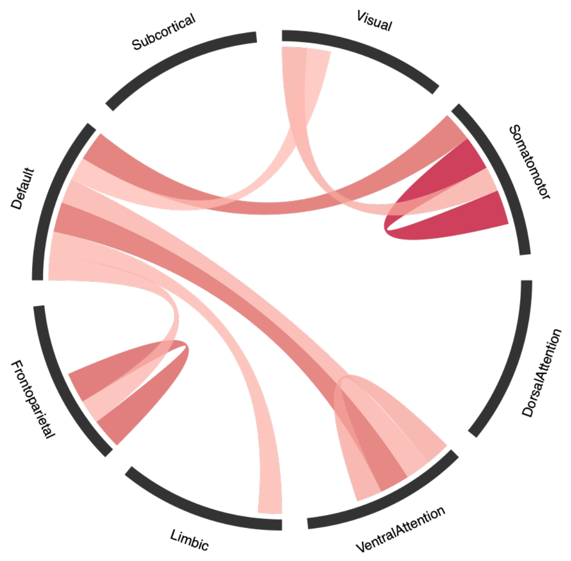


Figure G6

*Top 10 Reattributed (Yeo) Predictive Nodes: Structural Connectivity, Gordon Atlas*

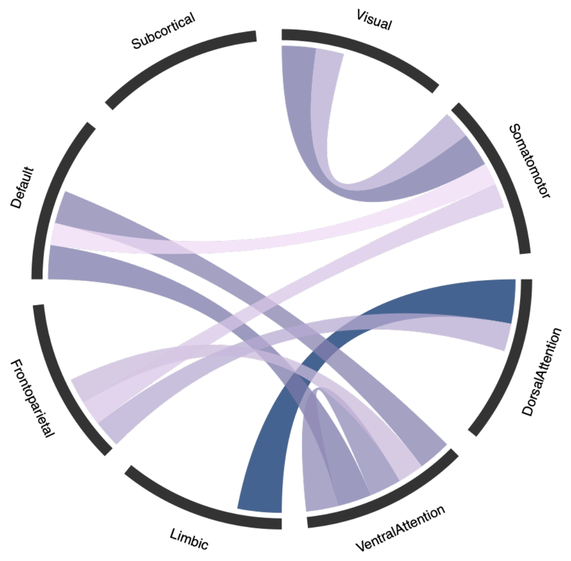


Figure G7

*Top 10 Reattributed (Yeo) Predictive Nodes: Functional (red) + Structural (blue) Connectivity, Brainnetome Atlas*

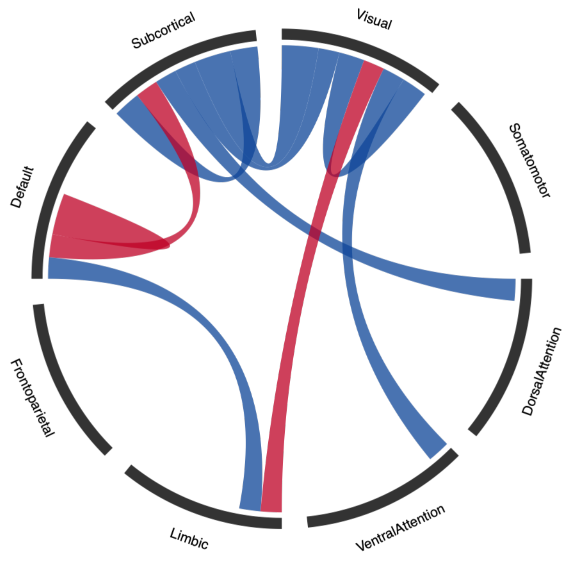


Figure G8

*Top 10 Reattributed (Yeo) Predictive Nodes: Functional (red) + Structural (blue) Connectivity, HCP MMP1 Atlas*

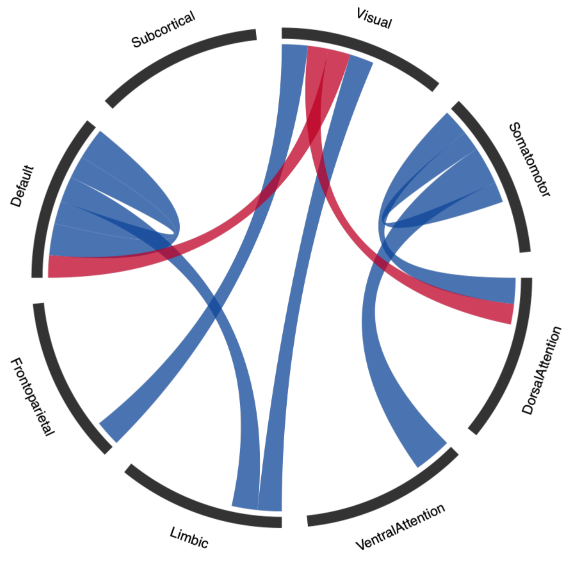


Figure G9

*Top 10 Reattributed (Yeo) Predictive Nodes: Functional (red) + Structural (blue) Connectivity, Gordon Atlas*

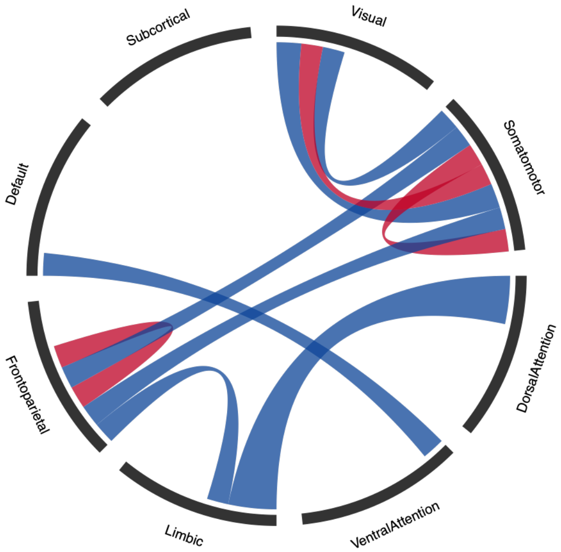

**Supplemental Material H (SM.H)**
**Fold Enrichment Clarification for Structural and Concatenated Dataset**

Figure H1

Structural Connectivity: Cross-Atlas Overlap of Top Yeo-7 Edges

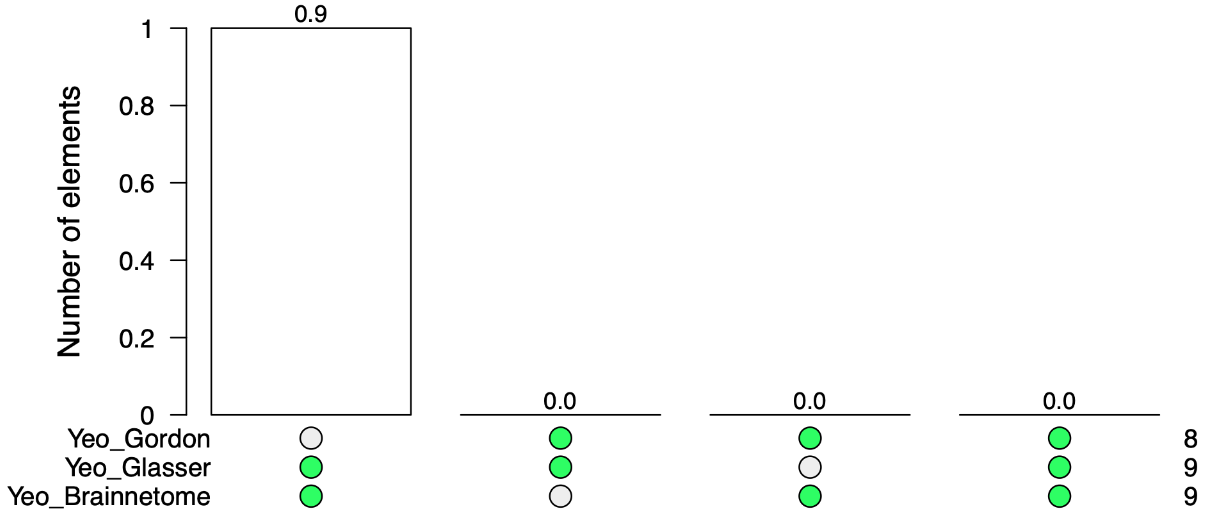


Figure H1

Concatenated Connectivity: Cross-Atlas Overlap of Top Yeo-7 Edges

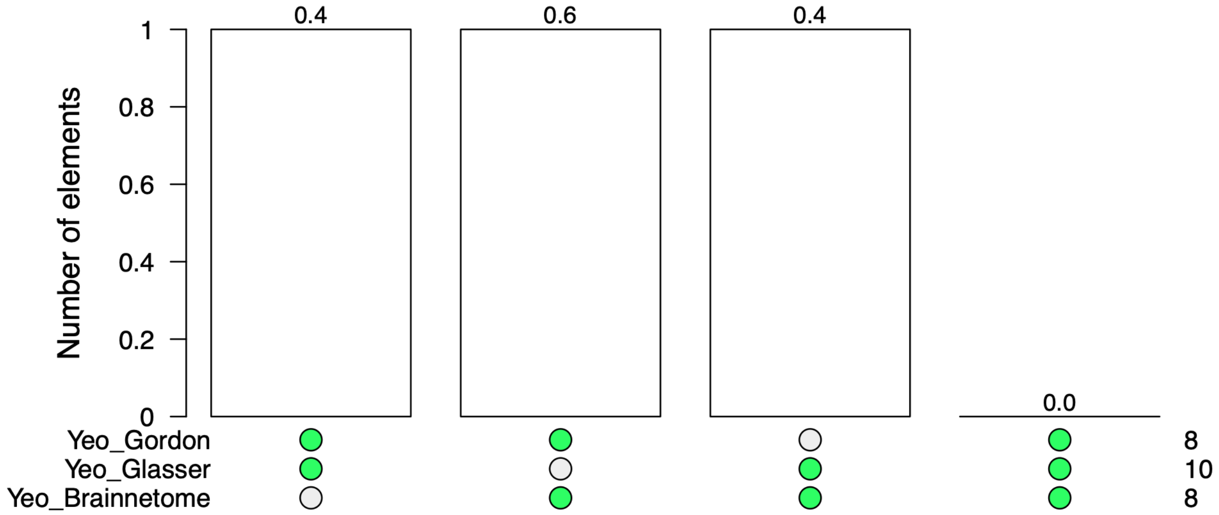
